# Functional mapping of synaptotagmin 1 Ca^2+^ ligands that mediate excitation-secretion coupling in neurons

**DOI:** 10.64898/2026.11.12.766256

**Authors:** Shweta Mishra, Nikunj Mehta, Matthew J. Dominguez, Emily R. Beckett, Mitch Wozney, Alen Vukovic, R. Bryan Sutton, Edwin R. Chapman

## Abstract

Synaptotagmin (SYT) 1 is a Ca^2+^ sensor that triggers rapid synaptic vesicle exocytosis, but how it is activated by Ca^2+^ remains enigmatic. SYT1 contains two Ca^2+^-sensing motifs, C2A and C2B. Structural studies have proposed that five acidic residues in each C2 domain mediate Ca^2+^-binding, but their functional relevance remains unclear. Here, we mutated each of the ten putative ligands and conducted side-by-side comparisons of Ca^2+^-binding activity and evoked glutamate release. Remarkably, all ten putative ligands are crucial for evoked synchronous release, overturning the view that binding of Ca^2+^ to C2A is dispensable. Importantly, Ca^2+^ binding to C2B is essential for synchronization. Furthermore, we report that two new human mutations, corresponding to Ca^2+^ ligands in C2B, are dominant-negative. Finally, four new crystal structures reveal how the choice of neutralizing substitution differentially alters SYT1 structure and, consequently, function.

## Introduction

Since the discovery of the first neurotransmitter in the early 20^th^ century^1,2^ the molecular mechanism(s) underlying the storage and release of these crucial chemical messengers garnered considerable attention. Subsequent studies established that neurotransmitters are stored in specialized secretory organelles, synaptic vesicles (SV), and are released via Ca^2+^-triggered exocytosis^3,4^. These advances prompted efforts to understand how Ca^2+^ signals are transduced to membrane fusion reactions, a process termed excitation-secretion coupling.

Progress on this front was made in the early 1980s, when a 65-kDa protein, p65, was discovered as a constituent of SVs and dense-core vesicles^5^. p65 was subsequently cloned and renamed synaptotagmin (SYT) 1^6^. From the sequence, the overall structure of SYT1 was predicted: it is an integral membrane protein with a cytoplasmic domain that harbors tandem C2 domains, C2A and C2B, that are connected by a short linker (Fig. 1a). These C2 domains are homologous to the second conserved (C2) motif first identified in isoforms of protein kinase C that are activated by Ca^2+^ and phosphatidylserine (PS)^7^. This observation suggested that SYT1 might be a Ca^2+^ sensor on the SV surface that can reversibly interact with anionic lipids, and these biochemical properties were directly demonstrated by Brose et al. in 1992^8^. Subsequent studies demonstrated that SYT1 is crucial for rapid and efficient synchronous SV exocytosis^9–13^ and chemical genetic experiments established that SYT1 functions as a divalent cation sensor that triggers synchronous neurotransmitter release^14^. Given the central role of SYT1 in coupling Ca^2+^ influx to release, it is perhaps unsurprising that missense mutations in *syt1* have been associated with neurodevelopmental disorders characterized by intellectual disability, motor delay, and abnormal movements, broadly referred to as Baker-Gordon syndrome^15–17^.

**Fig. 1:**
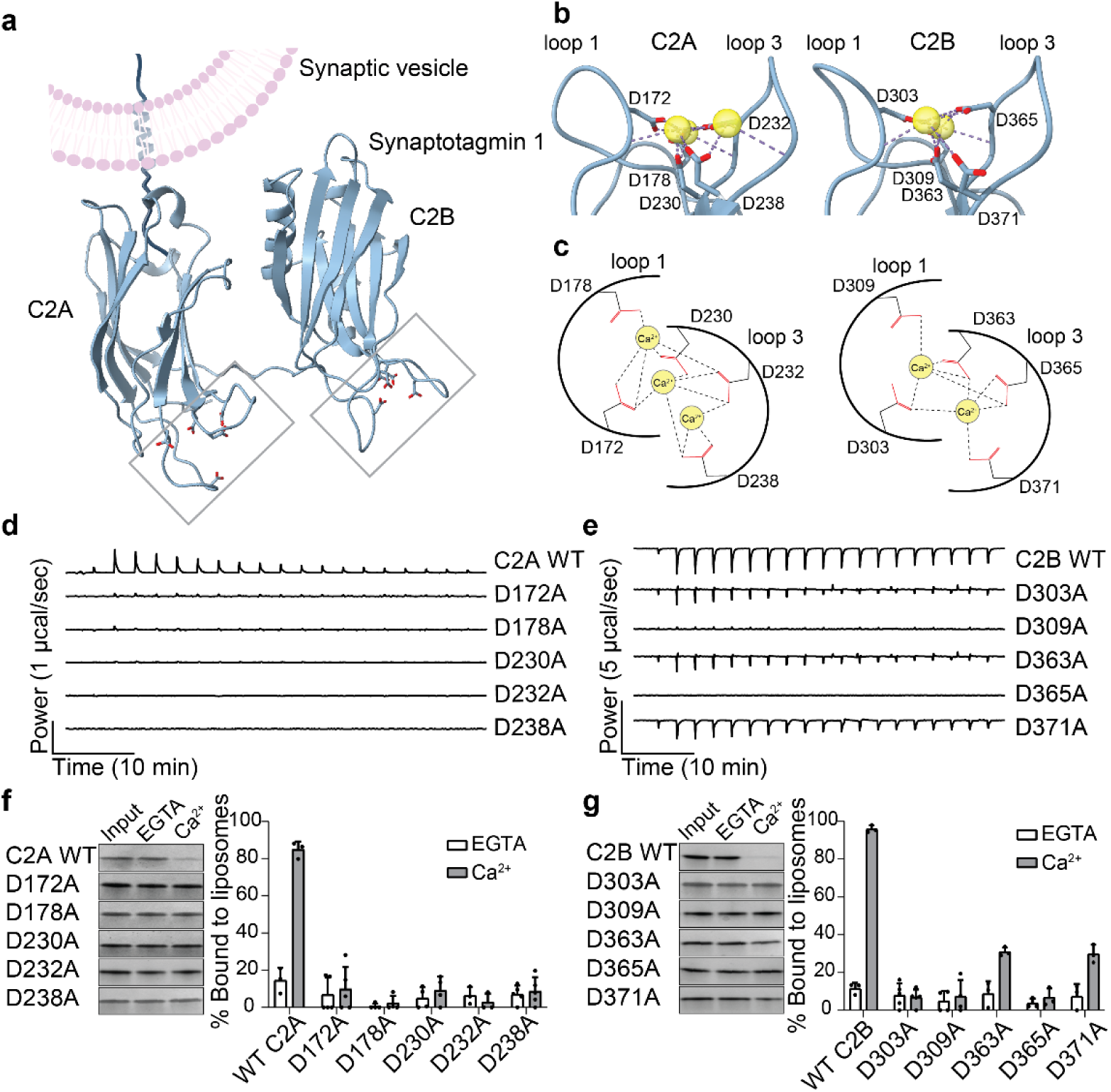
Neutralization of each putative Ca^2+^ ligand in the isolated C2 domains of SYT1 impairs Ca^2+^ and lipid binding activity. **a,** Schematic of full-length SYT1 spanning the synaptic vesicle (SV) membrane. Each C2 domain was rendered from crystal structures using ChimeraX^51^ (C2A: PDB 4WEE, C2B: PDB 6TZ3); the SV membrane and N-terminal segment were drawn using BioRender, the linker between C2A and C2B was drawn using Adobe Illustrator. Gray boxes highlight Ca^2+^-binding loops in each C2 domain; the Asp side chain oxygens are shown as red sticks. **b,c,** Enlarged image and schematic showing Ca^2+^-binding loops 1 and 3 in the indicated C2 domain; Ca^2+^ ions are shown as yellow spheres. Putative Ca^2+^ coordination lines (dashed), from crystal structures (C2A: PDB 7UFQ chain A, C2B: PDB 7TX9), were added using ChimeraX^51^. Representative isothermal titration calorimetry (ITC) traces showing the heat of Ca^2+^ binding to WT and calcium ligand mutant (CLM) forms of the isolated SYT1 C2A (**d**) and C2B domains (**e**); N = 3. Dissociation constants are reported in Supplementary Table 1. **f,** Left: Representative SDS-PAGE gels showing the supernatant from WT and CLM C2A-domain•liposome co-sedimentation experiments, in the absence (EGTA) and presence of Ca^2+^. Input refers to the protein-only control. Right: Quantification of % protein bound to liposomes, in the absence and presence of Ca^2+^; N ≥ 3. **g,** Same as panel (f), but for WT and CLM forms of C2B; N ≥ 3. Liposome binding of WT C2A and C2B, in the presence of Ca^2+^, exceeded that of every mutant in every replicate (Mann-Whitney U-test).

The first direct insight into how SYT1 senses Ca^2+^ was obtained when the crystal structure of the C2A domain was determined, revealing a novel Ca^2+^-binding pocket formed by two flexible loops, 1 and 3 (Fig. 1b), that protrude from one end of a rigid β-sandwich structure^18^. From this structure, along with subsequent NMR studies^19,20^, a model was generated in which the acidic side chains of five Asp residues in C2A (D172, D178, D230, D232, and D238 in the mouse) serve as ligands that coordinate potentially three Ca^2+^ ions (Fig. 1c). This work was then expanded to the C2B domain, where five homologous Asp residues (D303, D309, D363, D365, and D371) were proposed to coordinate the binding of two Ca^2+^ ions^21^. Notably, these Asp residues are largely conserved across species (Fig. S1). Upon binding Ca^2+^, the distal tips of loops 1 and 3 in each C2-domain partially penetrate lipid bilayers^22–24^ and this has been shown to play a key role in triggering exocytosis^14,25,26^.

Despite the identification of these putative Ca^2+^-binding residues more than thirty years ago, a consensus regarding their function has not been reached. Early studies of synaptic transmission at the *Drosophila* neuromuscular junction (NMJ) concluded that Ca^2+^ binding to the C2A domain was completely dispensable for function^27^. Consistent with this, the first rescue study using cultured mouse *syt1* KO neurons found that neutralization of D232 and D238 with Asn had no effect on transmission^28^. However, when Stevens and Sullivan examined the D232N mutant, they reported a gain-of-function^29^. This was followed by a report in which D232N was gain-of-function, whereas D238N was loss-of-function^30^. Yet another study focused on putative Ca^2+^ ligand mutations in C2A reported loss of function, further clouding the issue^31^. At present, the role of Ca^2+^ binding to the C2A domain remains unclear.

In contrast to the C2A domain, studies of putative Ca^2+^ ligand mutations in the C2B domain have been consistently found to disrupt excitation-secretion coupling in both the *Drosophila* NMJ and cultured mouse neurons^32,33^; it is generally accepted that C2B must bind Ca^2+^ for SYT1 to trigger synchronous release. The only study to systematically examine each of the five ligands in C2B concluded that the first, second, and third ligands were crucial for function, while the fourth and fifth ligands were dispensable^33^. However, this study was limited to autaptic cultures of hippocampal neurons, which can yield phenotypes that differ from those of brain slice preparations and mass-dissociated cultures^34,35^.

Another potentially confounding factor in the controversy regarding the putative acidic Ca^2+^ ligands in SYT1 concerns the choice of amino acid residues used to neutralize them. Past studies have indiscriminately used both Ala and Asn. We note that the Ala side chain is apolar but is smaller; since the loops are flexible, this was presumed to be well tolerated. In contrast, the Asn side chain preserves the pocket size but is polar and hence might contribute to interactions with Ca^2+^. To date, direct comparisons between substitutions with Asn versus Ala have not been reported.

A clear view of how the C2 domains of SYT1 bind Ca^2+^ to trigger exocytosis has yet to be obtained. To help resolve this issue, we neutralized each of the ten putative acidic ligands via substitution with Ala. Then, in side-by-side comparisons, we rigorously determined the effects of these mutations on Ca^2+^-binding activity *in vitro* and on SV exocytosis in cultured hippocampal neurons. In some cases, we also compared Ala and Asn substitutions and found that the choice of neutralizing residue can lead to profound differences in function, which we explored using X-ray crystallography. Furthermore, our analyses included two disease-associated missense mutations at the same putative Ca^2+^ ligand position in the C2B domain; we report that these mutations are dominant-negative. Contrary to previous studies, Ca^2+^ binding to both C2 domains is essential for SYT1 function; all ten acidic residues contribute to excitation-secretion coupling, but the two C2 domains play different roles in this process.

## Results

### Biochemical analysis of all ten putative Ca^2+^ ligands in SYT1

While several studies have examined the impact of putative Ca^2+^ ligand mutations in SYT1, a comprehensive side-by-side analysis of all ten ligands has yet to be carried out. We began to address the role of each proposed ligand in binding Ca^2+^ via mutagenesis and isothermal titration calorimetry (ITC), which directly measures the intrinsic Ca^2+^ affinity. Because Ca^2+^ binding is endothermic for C2A but exothermic for C2B, their enthalpies cancel in the intact cytoplasmic domain^36,37^; so we used the isolated C2 domains. Each Ca^2+^ injection is indicated by a peak in the thermogram, representing the enthalpy change upon Ca^2+^ binding to the indicated C2 domain (Fig. 1d,e). For the C2A domain, disruption of each individual Asp residue completely abolished Ca^2+^-binding activity of all three sites (Fig. 1d). In contrast, three of the Asp residues in the C2B domain – D303A, D363A, and D371A – exhibited residual, low-affinity Ca^2+^-binding activity (Fig. 1e), all less than that of WT C2B (Table S1). Interestingly, fitting of these ITC traces suggests that all three mutants retained both Ca^2+^ binding sites in C2B.

SYT1 triggers exocytosis, at least in part, by penetrating membranes ─ that harbor anionic phospholipids ─ in response to Ca^2+^ influx^14,25,26^. Importantly, acidic phospholipids increase the affinity of SYT1 for Ca^2+ 8^, so it remained possible that the SYT1 Ca^2+^ ligand mutants (CLMs) could still bind Ca^2+^ in the presence of membranes. However, adding liposomes bearing anionic lipids to the ITC experiments complicates interpretation because multiple interactions occur, including membrane binding, liposome aggregation, and oligomerization^38^. So, we analyzed each mutant using a well-characterized protein-liposome co-sedimentation assay^36^. Isolated C2 domains were mixed with liposomes (20% DOPS / 80% DOPC) in 0.2 mM EGTA or 0.5 mM Ca^2+^. Liposomes were collected by ultracentrifugation, and the supernatant was run on SDS-PAGE gels (Figs. 1f,g left); depletion of the supernatant reports binding (Figs. 1f,g, right). The WT C2A and C2B domains efficiently bound to membranes in the presence of Ca^2+^, and this activity was completely disrupted in all five C2A CLMs. In the case of C2B, two mutants – D363A and D371A – retained some degree (∼30%) of Ca^2+^-dependent liposome binding activity. These findings are consistent with the residual intrinsic Ca^2+^-binding activity of these two mutants observed in the ITC experiments. Interestingly, while D303A also had residual Ca^2+^ binding activity by ITC, this mutant failed to bind liposomes in response to Ca^2+^.

### Impact of putative C2A Ca^2+^ ligand mutations on synchronous glutamate release

Next, we determined the impact of each C2A Ca^2+^ ligand mutation on SV exocytosis in cultured mouse hippocampal neurons. WT SYT1 (termed SYT1 rescue) or C2A CLMs were expressed in *syt1* cKO neurons (hereafter termed *syt1* KO) using lentiviral transduction. Here and in all subsequent experiments, expression levels were titrated to match endogenous SYT1 levels (Fig. 2a). Co-staining with synaptophysin (SYP) showed that all mutants were correctly targeted to SVs (Fig. S2). To monitor evoked glutamate release, an optical glutamate sensor, SF-iGluSnFR.A184V, was co-expressed using a separate lentivirus ^39^. We carried out these initial experiments at a high [Ca^2+^]_e_ (10 mM) to avoid false negatives in release measurements. Raw traces (Fig. 2b, left) and histograms generated by binning the iGluSnFR peaks versus time (Fig. 2b, right), confirm the loss of synchronous release in *syt1* KO neurons that is rescued by WT SYT1. Averaged iGluSnFR traces (Fig. 2c) and comparisons of peak synchronous ΔF/F_0_ (Fig. 2d) reveal a complete loss of evoked synchronous glutamate release for the D178A, D230A, and D232A mutants, whereas there is a partial, weak rescue with the D172A and D238A mutants. These findings indicate that while each putative Ca^2+^- binding residue in C2A plays a role in triggering exocytosis, they do not function equally, suggesting that Ca^2+^ might bind C2A sequentially. Notably, and despite the fact that all C2A Ca^2+^ ligand mutations reduced the release of glutamate, they still restored the loss of synchronicity observed in the KO (Fig. 2e), consistent with the idea that C2B drives synchronization^40^.

**Fig. 2:**
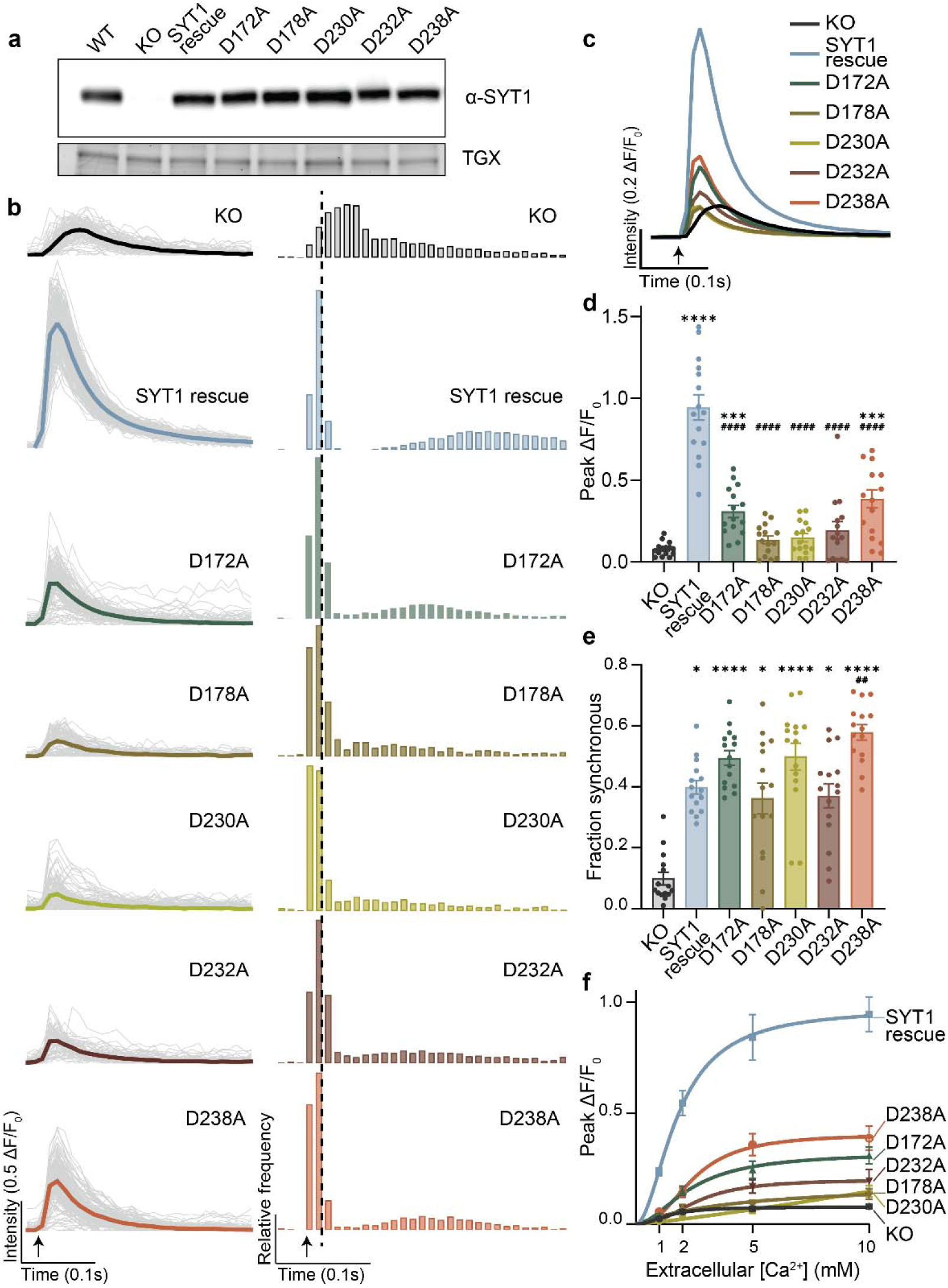
C2A calcium-ligand mutations decrease evoked glutamate release but preserve synchronicity. **a,** Representative anti-SYT1 immunoblot from cultured mouse hippocampal neurons: WT, *syt1* KO and *syt1* KOs expressing WT SYT1, as well as each of the C2A CLMs (D172A, D178A, D230A, D232A, and D238A), matched to endogenous levels, are shown. TGX serves as a loading control in this and all subsequent immunoblots. **b,** Left: Intensity-based glutamate-sensing fluorescent reporter (iGluSnFR) traces from representative fields of view (FOVs) showing individual regions of interest (ROIs; gray), and averaged (color-coded per condition) iGluSnFR signals (ΔF/F_0_), in 10 mM extracellular Ca^2+^ ([Ca^2+^]_e_); 300 ms epochs are shown. Right: Histograms of iGluSnFR peaks (10 ms bins). Averaged normalized traces **c,** and quantification of peak iGluSnFR ΔF/F_0_, **d,** for all conditions; each data point corresponds to a single FOV, n = 15 FOVs, N = 3 independent trials; error bars indicate SEM. Statistical significance was determined using Brown-Forsythe ANOVA test, with Dunnett’s T3 multiple comparisons test. **e,** Fraction of synchronous release measured as the ratio of the number of events in the first 20 ms after the stimulus over the total number of events 100 ms post-stimulus. Error bars represent SEM (Kruskal-Wallis test with Dunn’s multiple comparisons). **f,** Peak iGluSnFR ΔF/F_0_ for each C2A CLM at varying [Ca^2+^]_e_. Each point represents the mean of at least n = 14 FOVs at the indicated [Ca^2+^]_e,_ from N = 3 independent trials. Error bars indicate SEM; curves were fitted using the Hill equation. Statistical significance was determined using two-way ANOVA with Tukey’s multiple comparisons test. Detailed analyses are provided in the source data set. For all panels, post hoc comparisons to *syt1* KO and SYT1 rescue are indicated with * and #, respectively; ****, #### *p* < 0.0001; *** *p* < 0.001; ## *p* < 0.01; * *p* < 0.05.

To further characterize the Ca^2+^ sensitivity of these mutants, and to gain insight into release at physiological [Ca^2+^]_e_, we measured evoked glutamate release at varying [Ca^2+^]_e_ (1-10 mM) (Fig. 2f). At near-physiological Ca^2+^ (i.e., 2 mM), every C2A CLM failed to rescue the KO phenotype (Fig. S3).

### Disparate effects of D232A and D232N mutations in C2A

Next, we focused on a controversial issue: what occurs when D232 is replaced with an Asn? As discussed in the Introduction, this substitution has led to a variety of contradictory conclusions^28–30^. Our alanine screening approach already included D232A, thus providing a unique opportunity to compare two substitutions – Ala and Asn – at this position.

From the ITC experiments (Fig. S4a), it is evident that the D232N mutant, in stark contrast to D232A, retains some degree of Ca^2+^-binding activity. Fitting of the ITC trace suggests that the D232N mutant retains only two binding sites (Table S1), as opposed to three in WT C2A, indicating that this residue is a bona fide Ca^2+^ ligand. However, D232N failed to bind PS-bearing liposomes in the presence of Ca^2+^ (Fig. S4c), suggesting that loss of one Ca^2+^ binding site is sufficient to disrupt membrane interactions.

We then examined the effect of the D232N mutation on glutamate release, as described above; expression levels were determined via immunoblot analysis (Fig. 3a). In stark contrast to D232A, evoked synchronous glutamate release was fully rescued in the D232N mutant (Fig. 3b), with the peak synchronous amplitude reaching the same level as the SYT1 rescue (Fig. 3c). Moreover, like D232A, D232N restored the synchronicity of release (Fig. 3d); D232N also conferred a similar Ca^2+^-sensitivity for release compared with SYT1 rescue (Fig. 3e). These results reveal that neutralization of D232 with either Ala or Asn can have profoundly different effects on C2A function; this sheds new light on the controversies surrounding the Ca^2+^ ligands of SYT1, as these two substitutions have been used somewhat interchangeably in previous studies.

**Fig. 3:**
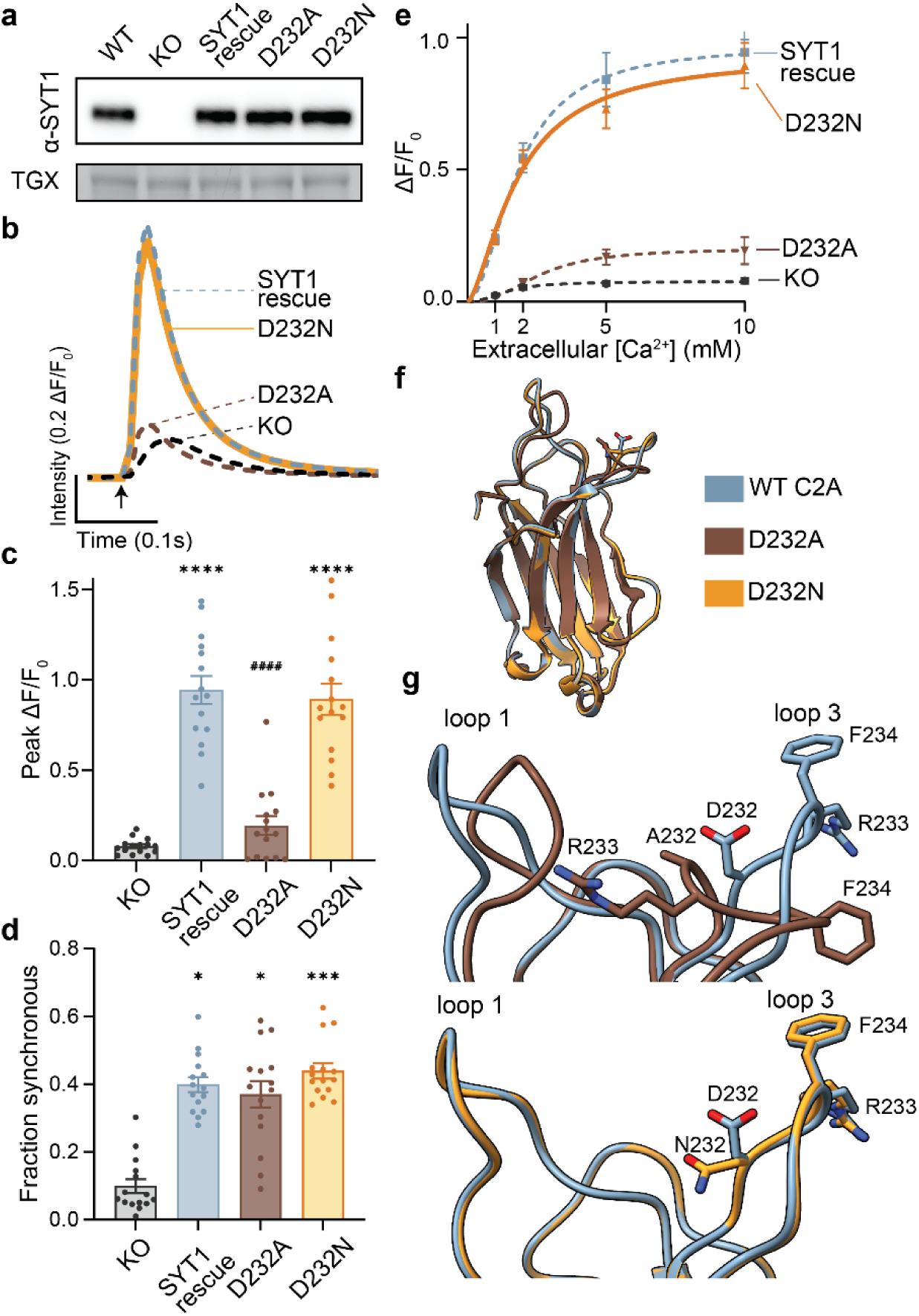
Choice of neutralizing substitution at position 232 has drastically different effects on glutamate release. **a,** Representative immunoblot from cultured mouse hippocampal neurons showing WT, *syt1* KO, and *syt1* KOs expressing endogenous levels of SYT1 rescue, D232A, and D232N mutants. Averaged normalized traces **b,** and quantification of peak iGluSnFR ΔF/F_0_, **c,** in 10 mM [Ca^2+^]_e;_ error bars indicate SEM (n = 15 FOVs, N = 3). For comparison, data for KO, SYT1 rescue, and D232A are replotted from Fig. 2 and shown as dashed lines. Statistical significance in (c) was determined using the Brown-Forsythe ANOVA test with Dunnett’s T3 multiple comparisons test. **d,** Fraction of synchronous release; error bars represent SEM (Kruskal-Wallis test with Dunn’s multiple comparisons) **e,** Peak iGluSnFR ΔF/F_0_ for at varying [Ca^2+^]_e_. Each point represents the mean of n = 15 FOVs at the indicated [Ca^2+^]_e,_ from N = 3 independent trials. Error bars indicate SEM; curves were fitted using the Hill equation. Statistical significance was determined using two-way ANOVA and Tukey’s multiple comparisons test. Detailed analyses are provided in the source data set. **f,** Superimposition of the asymmetric units of SYT1 C2A D232A (PDB: 9ZH6, chain B) and SYT1 C2A D232N (PDB: 9ZKH), compared with WT SYT1 C2A (PDB:4WEE). **g,** Magnified view of Ca^2+^ binding loops 1 and 3, with relevant residues labelled; top: SYT1 C2A D232A compared to WT SYT1 C2A, bottom: SYT1 C2A D232N compared to WT SYT1 C2A. For all panels, post hoc comparisons to *syt1* KO and SYT1 rescue are indicated with * and #, respectively; ****,#### *p* < 0.0001; *** *p* < 0.001; * *p* < 0.05

To address the differences between D232A and D232N, we undertook a structural approach using X-ray crystallography. We crystallized D232A and D232N at 2.74 and 1.20 Å resolution, respectively, and overlaid them with a previously published crystal structure of C2A WT^41^ (Fig. 3f). Both variants retained the overall C2-domain β-sandwich fold, indicating that the substitutions did not alter the global architecture; however, there were localized rearrangements discussed below. The largest differences were observed in D232A within Ca^2+^-binding loop 1 (E167–D178) and loop 3 (D230–D238), whereas D232N remained essentially unchanged relative to apo WT (Fig. S5a). However, normalized B-factors indicate that loop 1 is intrinsically mobile in WT, D232N, and D232A chain A, while the same region in D232A chain B is comparatively well ordered (Fig. S5b), likely due to crystal-packing interactions. Despite differences in loop 1 mobility between the two D232A molecules, both modeled loop 1 conformations more closely resembled the Ca^2+^-bound state of 7U4Q^41^ and the Pb^2+^-bound structure 5VFE^42^ than the apo WT conformation. Further, loop 3 in D232A adopted a distinct, well-ordered conformation (Fig. 3g, top) not observed in the apo or metal-bound reference structures; Arg233 penetrated the Ca^2+^-binding pocket, and Asp238 was rotated away from its Ca^2+^-coordinating position, rendering the observed structure incompatible with known C2-domain metal-binding conformations. Comparative electrostatic calculations indicated that the membrane-facing surface changed from negative in WT to positive in D232A in the absence of bound Ca^2+^ (Fig. S5c). The membrane-penetrating residue in loop 1, Met173^23^, remained solvent-exposed, but the membrane-penetrating residue in loop 3, Phe234c^22^ was redirected toward the β-sandwich and became substantially less accessible (Fig. 3g, top). This local rearrangement in D232A also extended into the β4 (189-194) -β5 (208-212) hydrophobic core (Fig. S5d). Phe193 on β-strand 4 and Phe210 and Phe212 on β-strand 5 adopted orientations that differed from those observed in previous apo and metal-bound WT structures. Phe193 and Phe210 showed conserved rotamer changes in both D232A molecules. The close Phe193-Phe212 contact, present in the WT and metal-bound reference structures, was lost. Phe193 instead remained closely packed against Phe210. Phe212 also changed orientation, although its precise rotamer differed between the two D232A molecules. D232A therefore reproduces the electrostatic component of the metal-bound state while disrupting canonical metal coordination and the membrane-facing geometry of loop 3, potentially explaining why this mutant is unable to rescue function.

In the structure of the D232N mutant domain, Asn232 mimics the rotamer change observed for Asp232 in the Ca^2+^-bound C2A structure (Fig. 3g, bottom, Fig. S5e). Arg233 remained directed away from the Ca^2+^-binding pocket. Met173 and Phe234 also retained outward-facing, solvent-exposed orientations. Asn232 formed a local interaction network with the Lys236 backbone carbonyl and Asp238, while its carbonyl oxygen remained directed toward the cation-binding site. Thus, D232N preserves the overall architecture of the WT Ca^2+^-binding region despite removal of the Asp232 carboxylate.

### Role of each putative Ca^2+^ ligand in C2B in excitation-secretion coupling

Having addressed the impact of Ca^2+^ ligand mutations in C2A, we next determined how analogous mutations in the C2B domain affect evoked synchronous glutamate release. All C2B mutants were expressed to match endogenous levels of SYT1 (Fig. 4a). As with C2A, none of the C2B CLMs affected the localization of SYT1, demonstrated by their co-localization with the SV marker SYP (Fig. S6). From the raw traces (Fig. 4b, left) and histograms of iGluSnFR peaks (Fig. 4b, right), it is evident that each C2B Ca^2+^ ligand mutation severely impaired synchronous release at high [Ca^2+^]_e_ (10 mM). Averaged iGluSnFR traces (Fig. 4c) and peak synchronous ΔF/F_0_ (Fig. 4d) reveal a complete loss of evoked release in D303A, D363A, and D365A, with D309A yielding even less glutamate release than *syt1* KO neurons. Interestingly, D371A was the only mutant that trended toward a weak, partial rescue. These findings contrast earlier work using autaptic cultures^33^ and support the idea that all five putative Ca^2+^ binding residues in C2B are crucial for SYT1 function.

**Fig. 4:**
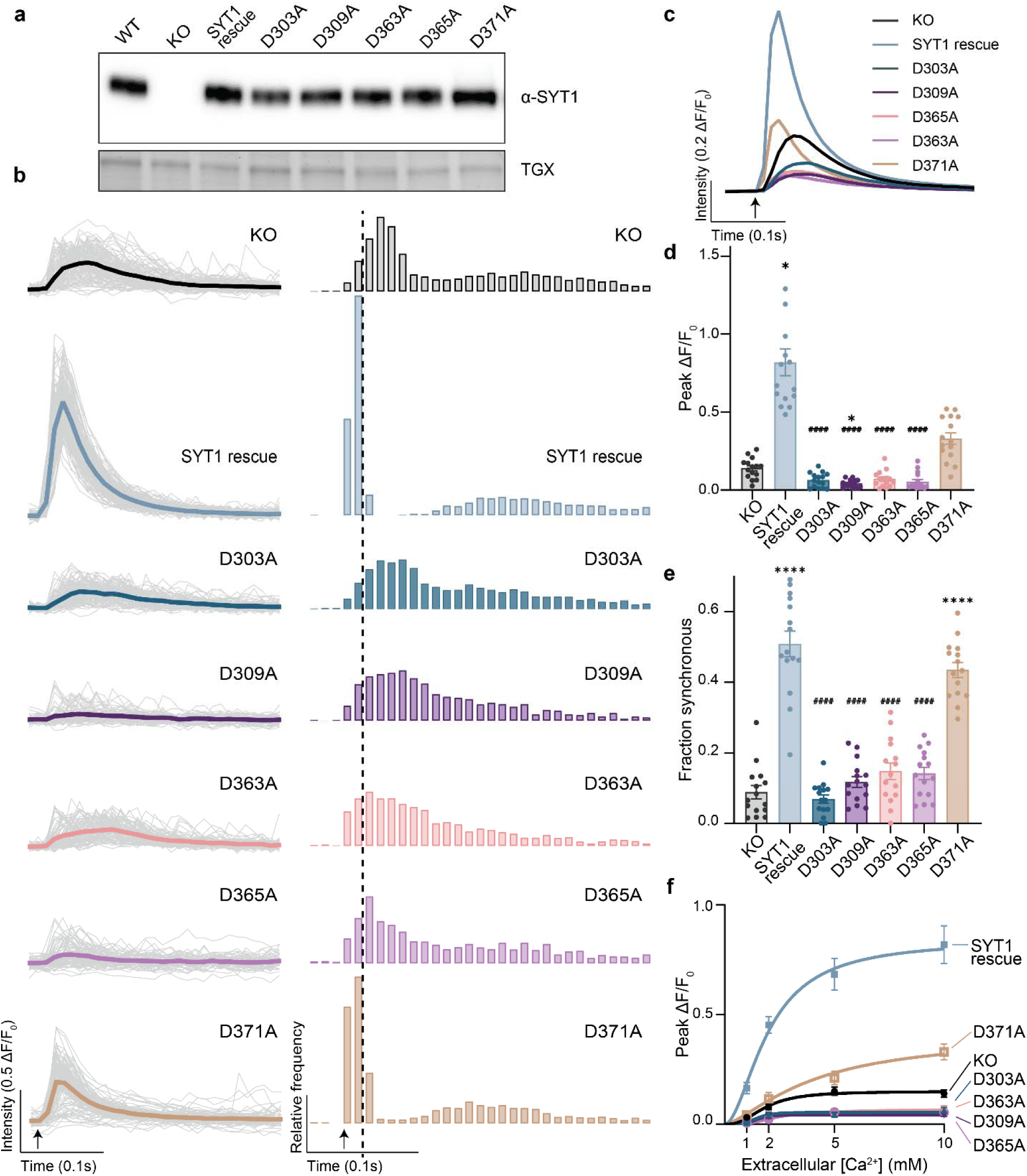
C2B calcium-ligand mutations decrease evoked glutamate release and disrupt synchronicity. **a,** Representative immunoblot from cultured mouse hippocampal neurons showing WT, *syt1* KO, and *syt1* KOs expressing SYT1 rescue and each of the C2B CLMs (D303A, D309A, D363A, D365A, and D371A), matched to endogenous SYT1 levels. **b,** Left: iGluSnFR traces from representative FOVs showing individual ROIs (gray) and averaged (color-coded per condition) iGluSnFR signals (ΔF/F_0_) in 10 mM [Ca^2+^]_e_; 300 ms epochs are shown. Right: Histograms of iGluSnFR peaks (10 ms bins). Averaged normalized traces **c,** and quantification of peak iGluSnFR ΔF/F_0_, **d,** for all conditions; error bars indicate SEM, n = 15 FOVs, N = 3 independent trials. Statistical significance was determined by Kruskal-Wallis test, with Dunn’s multiple comparisons. **e,** Fraction of synchronous release for all conditions in (c), with error bars representing SEM (Brown-Forsythe ANOVA test and Dunnett’s multiple comparisons test). **f,** Peak iGluSnFR ΔF/F_0_ for each C2B CLM at varying [Ca^2+^]_e_. Each point represents the mean of at least 15 FOVs at the indicated [Ca^2+^]_e,_ from at least 3 independent trials. Error bars indicate SEM; curves fitted using Hill equation. Statistical significance was determined using two-way ANOVA with Sidak’s multiple comparisons test. Detailed analyses are provided in the source data set. For all panels, post hoc comparisons to *syt1* KO and SYT1 rescue are indicated with * and #, respectively; ****,#### *p* < 0.0001; * *p* < 0.05

In contrast to each of the C2A CLMs, where the synchronicity of release was maintained, four of the five CLMs in C2B completely disrupted synchronization (Fig. 4e). These findings are congruent with the idea that the C2B domain of SYT1 is necessary and sufficient to synchronize release^40^. Importantly, the current study demonstrates that the Ca^2+^-binding activity of this domain underlies synchronization; clearly, C2B controls release kinetics ^14,40^. The lone exception was the D371A mutant, which still retains synchronicity. Ca^2+^ dose-response experiments show a complete loss of function across all C2B CLMs at physiological [Ca^2+^]_e_ (Fig. 4f; Fig. S7).

### Pathogenic SYT1 C2B mutations have little effect on structure but are dominant-negative for evoked glutamate release

Having shown that all ten proposed SYT1 ligands play roles in coordinating Ca^2+^, we next turned our attention to *de novo* heterozygous missense mutations in human *syt1* that are associated with severe neurodevelopmental disorders^15–17^. Most previously reported pathogenic SYT1 mutations cluster in the C2B domain^15–17^. Here, we explored two novel disease-associated variants at the same position in C2B: D309A and D309N (detailed clinical histories of the patients are provided in the Methods). D309A was already characterized in the preceding section; by comparing this mutation with D309N, we can determine whether there are functional differences between these mutants, analogous to the D232A-D232N comparisons above.

We examined the intrinsic properties of the D309N mutant using ITC and the isolated C2B domain. In contrast to D309A, where Ca^2+^ binding and co-sedimentation with liposomes were abolished, D309N retained Ca^2+^-binding activity, albeit with a reduced affinity at both binding sites (Fig. S4b and Table S1), as well as a partial ability to co-sediment with liposomes in response to Ca^2+^ (Fig. S4d).

The effect of the D309N mutant on function was determined after expression in *syt1 KO* neurons (Fig. 5a). Similar to D309A above, D309N abrogated evoked synchronous glutamate release at 10 mM [Ca^2+^]_e_ (Fig. 5b), yielding a peak synchronous amplitude that was even lower than in *syt1* KO neurons (Fig. 5c). In addition, all observed residual release was asynchronous (Fig. 5d). These findings held across varying [Ca^2+^]_e_ (Fig. 5e) and indicate a complete disruption of the Ca^2+^-sensitivity of release. These findings sharply contrast our observations regarding Asn versus Ala at position 232 in C2A, where Ala disrupted function while Asn was largely tolerated. So, substitution with Asn or Ala can either yield no difference or can have profound differences, depending on which Ca^2+^ ligand is being altered.

**Fig. 5:**
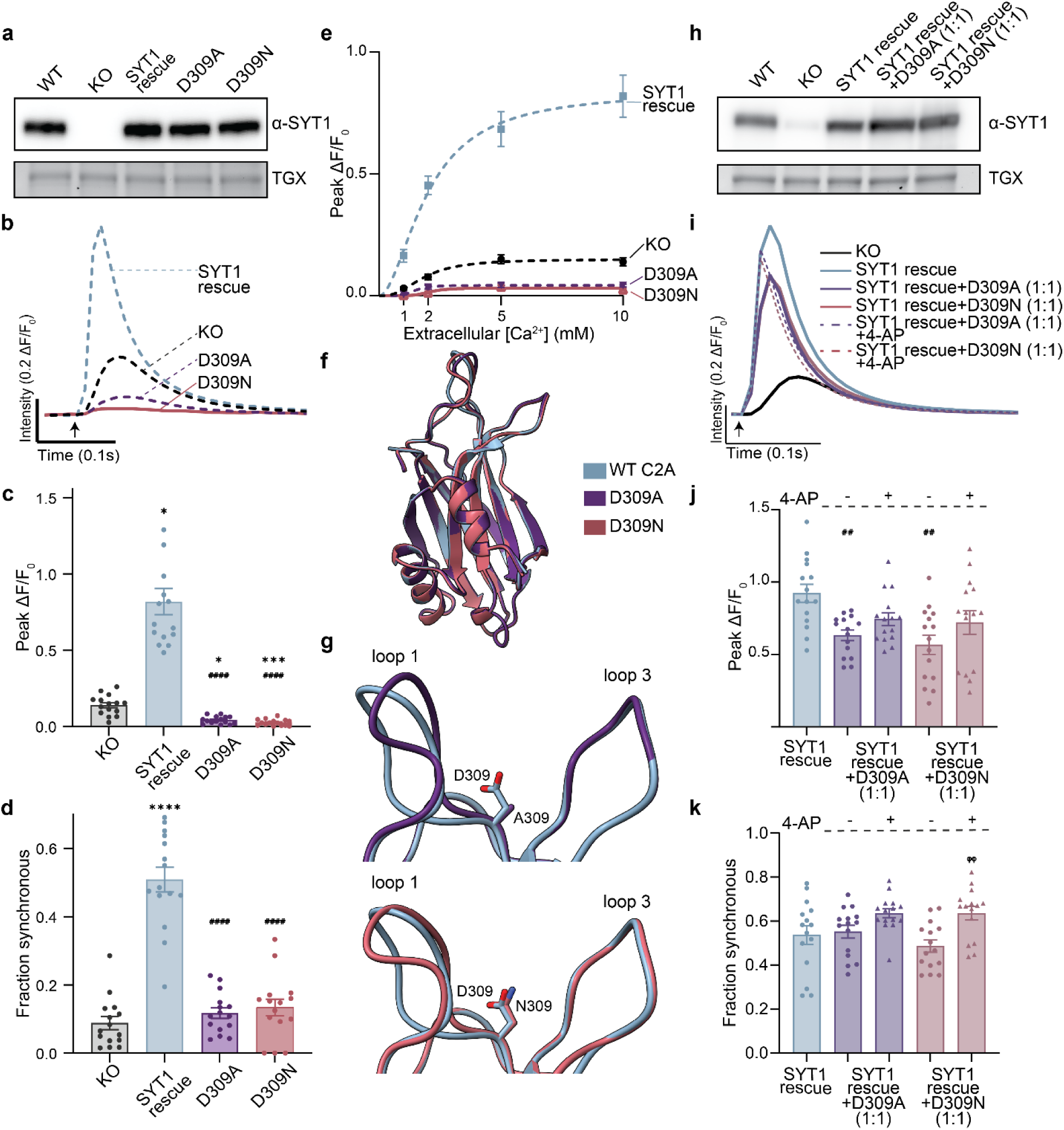
Disease-associated C2B calcium-ligand mutants exhibit a dominant-negative effect on evoked glutamate release but do not impair synchronicity. **a,** Representative anti-SYT1 immunoblot from cultured mouse hippocampal neurons showing WT, *syt1* KO, and *syt1* KOs expressing SYT1 rescue, D309A, and D309N mutants, all matched to endogenous SYT1 expression levels. Averaged normalized traces **b,** and quantification of peak iGluSnFR ΔF/F_0_. **c,** in 10 mM [Ca^2+^]_e;_ error bars indicate SEM, n = 15 FOVs, N = 3 independent trials. For comparison, data for KO, SYT1 rescue, and D309A are replotted from Fig. 4 as dashed lines; this also applies to panels (d), (e) and (f). Statistical significance in (c) was determined using Kruskal-Wallis test with Dunn’s multiple comparisons. **d,** Fraction of synchronous release, determined as in Fig. 2E; error bars represent SEM (Brown-Forsythe ANOVA test and Dunnett’s multiple comparisons test). **e,** Peak iGluSnFR ΔF/F_0_ as a function of [Ca^2+^]_e_. Each point represents the mean of at least 14 FOVs at the indicated [Ca^2+^]_e,_ from at least 3 independent trials. Error bars indicate SEM; curves were fitted using the Hill equation. Statistical significance was determined using two-way ANOVA and Sidak’s multiple comparisons test. Detailed analyses are provided in the source data set. **f,** Superimposition of the asymmetric units of SYT1 C2B D309A (PDB: 9ZJL) and SYT1 C2B D309N (PDB: 9ZF1), compared with WT SYT1 C2B (PDB:6TZ3). **g,** Magnified view of Ca^2+^ binding loops 1 and 3, with relevant residues labelled; top: SYT1 C2B D309A compared to WT SYT1 C2B, bottom: SYT1 C2B D309N compared to WT SYT 1 C2B. **h,** Representative anti-SYT1 immunoblot from cultured mouse hippocampal neurons showing WT, *syt1* KO, and *syt1* KOs expressing SYT1 rescue (matched to endogenous levels), SYT1 rescue + D309A, and SYT1 rescue + D309N in 1:1 ratio (∼2x of endogenous SYT1). **i,** Averaged normalized traces from each rescue condition shown in panel; n = 15 FOVs, N = 3 independent trials. **j,** Quantification of peak iGluSnFR ΔF/F_0_ (Brown-Forsythe ANOVA test with Dunnett’s T3 multiple comparisons test), and **k,** fraction of synchronous release (Brown-Forsythe ANOVA test with Dunnett’s T3 multiple comparisons test), before and after application of 5 μM 4-aminopyridine, from three independent trials in 2 mM [Ca^2+^]_e_, again for each condition. For all panels, post hoc comparisons to *syt1* KO and SYT1 rescue are indicated with * and #, respectively; denotes the comparison between before-and after-treatment conditions; ****,#### *p* < 0.0001; *** *p* < 0.001; ##, *p* <0.01; * *p* < 0.05

To further characterize the pathogenic SYT1 variants, we determined crystal structures of D309A and D309N to 1.07 and 1.05 Å resolution, respectively. Both C2B mutants retained the overall C2 domain β-sandwich fold and closely aligned with the apo WT SYT1 C2B structure (PDB 6TZ3) ^17^(Fig. 5f). The largest differences were observed within loop 1 (K297–P310), whereas the other Ca^2+^-binding loop, loop 3 (D363–D371), remained essentially unchanged (Fig. 5g, Fig. S8a). However, elevated normalized B-factors in WT and both mutants indicate that loop 1 is intrinsically mobile, with increased local disorder in the mutants, particularly D309A (Fig. S8b); hence, we do not interpret these modest differences as evidence for a distinct mutation-induced conformation. Further, we compared the mutant structures with the Ca^2+^-bound C2B structures 1TJX, 1UOV, and 7TX9 ^41,43^. Across these structures, D309 participated directly in Ca^2+^ coordination, with the side-chain carboxylate contributing at least one coordinating Ca^2+^ interaction, in addition to the intramolecular interaction with the Asn333 backbone in loop 2 (Fig. S8c, Supplementary Table 3). D309A removes the side-chain carboxylate entirely, leaving the Ca^2+^-binding cavity open and solvent accessible (Fig. S8d). D309N preserves a similarly sized side chain and retains a carbonyl oxygen that maintains the interaction with the Asn333 backbone (Fig. S8e). Thus, the pathogenic mutations at D309 disrupt a direct Ca^2+^ ligand without substantially reorganizing the surrounding binding-site architecture.

In all these experiments, disease-associated mutations were expressed in a *syt1* KO background. However, in patients, these mutations are heterozygous. To mimic these conditions and assay for dominant-negative activity ^17^ we co-expressed the mutants, in a *syt1* KO background, with WT SYT1 at a 1:1 ratio (Fig. 5h). As described previously ^17^ we also used 4-Aminopyridine (4-AP), an FDA-approved K^+^-channel antagonist, to enhance presynaptic Ca^2+^ influx and potentially mitigate any dominant-negative effects caused by the mutations. Averaged traces (Fig. 5i) and peak synchronous amplitudes (Fig. 5j) at 2 mM [Ca^2+^]_e_ show that both mutants are dominant-negative, with ∼30% and 40% reductions in peak synchronous release for D309A and D309N, respectively. Treatment with 5 μM 4-AP slightly increased the extent of synchronous release, but this did not reach statistical significance under these conditions. These observations warrant further study, as 4-AP was effective in a previous report examining different pathological C2B mutants ^17^. Interestingly, synchronicity ─ under these dominant-negative conditions ─ remained unaffected (Fig. 5k) and was further enhanced by treatment with 4-AP; this latter effect reached significance for D309N (Fig. 5k).

### Neuralization of all five Ca^2+^ ligands in C2A partially suppresses the loss-of-function phenotype of single-point mutants

Throughout this study, we mutated individual putative Ca^2+^-binding residues; for completeness, we simultaneously mutated all five residues in C2A or C2B, expecting this to completely abolish function. Indeed, Ca^2+^ and membrane-binding activity were abolished in both C2A_5A_ and C2B_5A_ mutants (Fig. S4). We then expressed these mutants in *syt1* KO neurons (Fig. 6a) and assessed their impact on glutamate release as detailed in previous sections. From the raw traces (Fig. 6b, left) and histograms of iGluSnFR peaks (Fig. 6b, right) at 2 mM [Ca^2+^]_e_, it is evident that C2B_5A_ disrupts all evoked glutamate release. Unexpectedly, as shown in the averaged iGluSnFR traces (Fig. 6c) and the peak synchronous amplitude plot (Fig. 6d), C2A_5A_ retains ∼70% of synchronous release relative to the SYT1 rescue. Notwithstanding, the fraction of synchronous release in both C2A_5A_ and C2B_5A_ (Fig. 6e) aligns with our observations from the previous sections: Ca^2+^ binding to C2B is necessary to synchronize evoked glutamate release, whereas both C2 domains are required to fully rescue the extent of synchronous release^40^.

**Fig. 6:**
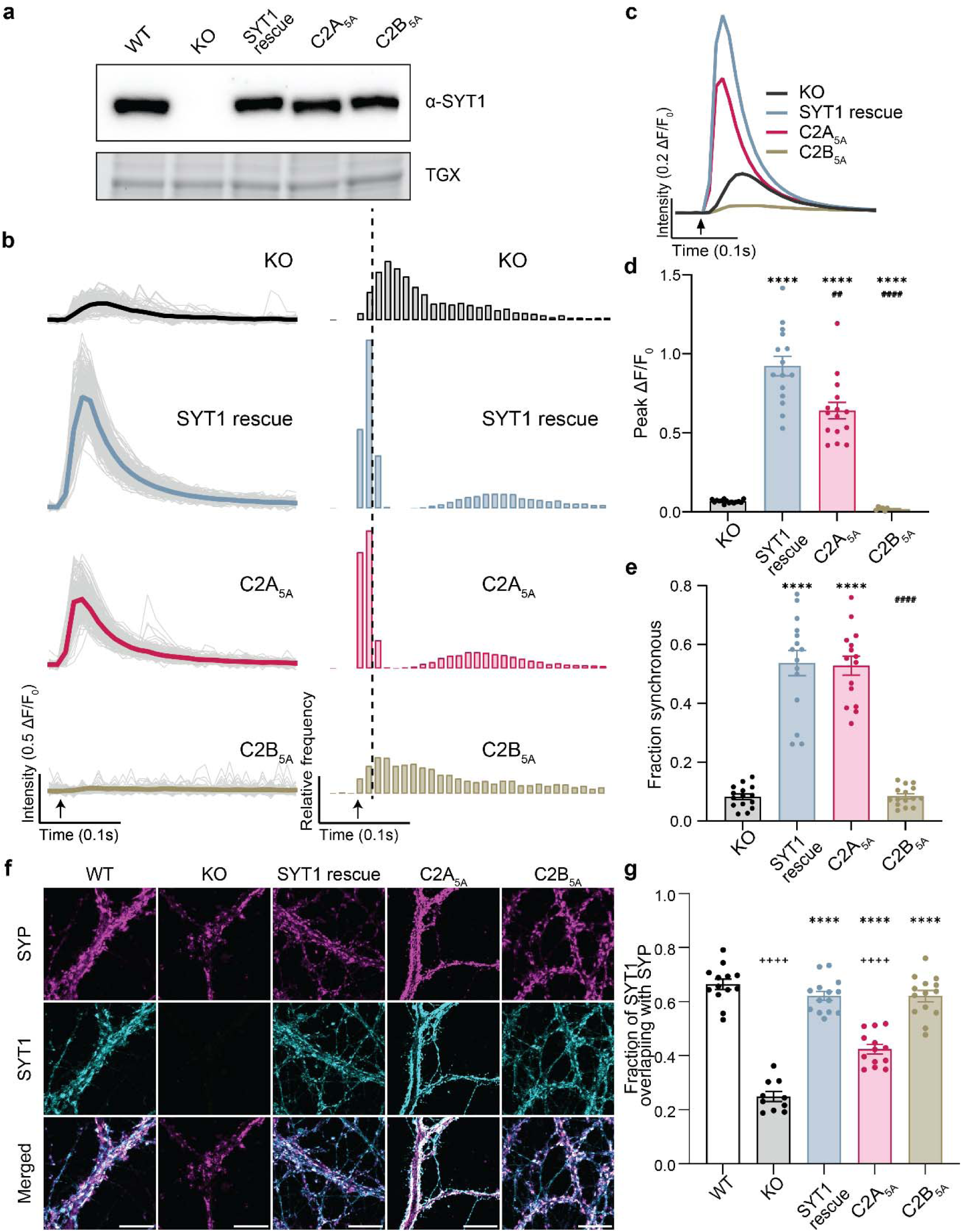
Concurrent neutralization of all five putative calcium ligands results in mis-localization and partial function in C2A, but not in C2B. **a,** Representative anti-SYT1 immunoblot from cultured mouse hippocampal neurons showing WT, *syt1* KO, and *syt1* KOs expressing SYT1 rescue, C2A_5A_, and C2B_5A_ mutants, all matched to endogenous SYT1 expression levels. **b,** Left: iGluSnFR traces from representative FOVs showing individual ROIs (gray) and averaged (color-coded per condition) iGluSnFR signals (ΔF/F_0_) in 2 mM [Ca^2+^]_e_; 300 ms epochs are shown. Right: Histograms of iGluSnFR peaks (10 ms bins). **c,** Averaged normalized traces and **d,** quantification of peak iGluSnFR ΔF/F_0;_ n =15 FOVs, N =3 independent trials; error bars indicate SEM (Brown-Forsythe ANOVA test with Dunnett’s T3 multiple comparisons test). **e,** Fraction of synchronous release; error bars represent SEM (Brown-Forsythe ANOVA test with Dunnett’s T3 multiple comparisons test). **f,** Representative super-resolution fluorescence images of mouse hippocampal neurons. Images of WT, *syt1* KO, and *syt1* KOs expressing WT SYT1 rescue, C2A_5A_, and C2B_5A_, immunolabeled for presynaptic marker synaptophysin (SYP) and SYT1; in the last row, both signals were merged to show co-localization. Scale bar 10 μm. **g,** Quantification of the colocalization of SYT1 in each condition with SYP measured using Manders’ overlap coefficients. Each data point corresponds to a single FOV; n = 10-14 FOVs, N = 2 independent trials. Statistical significance was determined using the Brown-Forsythe ANOVA test, with Dunnett’s T3 multiple comparisons test. For all panels, post hoc comparisons to WT, *syt1* KO and SYT1 rescue are indicated with +, * and #, respectively; ****, ####, ++++ *p* < 0.0001; ## *p* < 0.01

The inability of C2B_5A_ to rescue transmission is not due to a protein-sorting defect, as this mutant was properly targeted to SVs (Fig. 6f,g; Fig. S9). This was the expected result, as mutations in the Ca^2+^ and membrane-binding loops of SYT1 have consistently been reported not to affect synaptic targeting, perhaps because these loops are flexible. However, the C2A_5A_ mutant broke this rule, as it was largely localized extrasynaptically at the plasma membrane. After biosynthesis and transport to axons, SYT1 is initially delivered to the plasma membrane and is subsequently incorporated into SVs, presumably via endocytosis^44^. Together, these findings suggest that the C2A_5A_ mutant fails to become internalized.

Regardless of the unexpected findings regarding C2A_5A_, the observations reported here provide strong evidence that each individual Ca^2+^ ligand in C2A and in C2B serves as a bona fide Ca^2+^-binding residue, and that both C2 domains are required for efficient Ca^2+^-triggered SV exocytosis, with C2B also synchronizing release.

## Discussion

In the current study, we addressed the unresolved question concerning which of the ten putative acidic Ca^2+^ ligands in SYT1 ─ proposed from structural studies ─ coordinate Ca^2+^ ions and underlie its role as a Ca^2+^ sensor for synchronized neurotransmitter release. Numerous biochemical and functional studies, over the past few decades, have reported conflicting findings regarding the importance of these residues, and a coherent view has yet to emerge. Indeed, some studies concluded that Ca^2+^ binding to C2A was dispensable for function^27,28^. Here, we systematically mutated each putative Ca^2+^ ligand in each C2 domain and, side by side, assayed Ca^2+^ binding activity *in vitro* and excitation-secretion coupling in cultured neurons.

In our biochemical assays with isolated C2A, all five single D-to-A mutants completely disrupted the low-affinity, intrinsic Ca^2+^-binding activity of this domain. Since anionic lipids increase the affinity of SYT1 for Ca^2+^, we also conducted Ca^2+^-dependent lipid-binding assays and found that this activity was abolished. Importantly, each of the five putative Ca^2+^ ligand mutations in C2A completely disrupted the ability of SYT1 to rescue evoked synchronous glutamate release at physiological [Ca^2+^]_e_. Interestingly, two mutants ─ D172A and D238A – supported low levels of release at high [Ca^2+^]_e_ (i.e., 10 mM). These findings suggest that the other residues, D178, D230, and D232, might bind Ca^2+^ first. The residual function of the D172A and D238A mutants also likely involves the intact adjacent C2B domain, a point we return to below. A second crucial point is that while the CLMs failed to rescue the amplitude of evoked responses, each mutant was able to synchronize the residual release in the KOs, likely due to the presence of the neighboring C2B domain^40^.

In the literature, D232 has been a particularly confusing position, perhaps in part because it has been neutralized by substitution with either Ala or Asn. To address this, we directly compared these two mutations. Unlike C2A D232A, C2A D232N exhibited some degree of Ca^2+^ binding activity by ITC but failed to bind membranes in response to Ca^2+^. Surprisingly, the D232N mutant fully rescued evoked synchronous glutamate release in dissociated neurons. These unexpected results suggest that the residual weak Ca^2+^ binding sites in D232N might be “permissive”, allowing the adjacent C2B domain to drive membrane penetration of the nearby mutant C2A domain by ‘dragging’ its membrane-penetrating loops into the plasma membrane^45^. The same idea might apply to the D172A and D238A mutants, to yield a low degree of excitation-secretion coupling at high [Ca^2+^]_e_.

To better understand why Ala and Asn have such dramatically different effects at position 232 in C2A, we determined the crystal structures of each mutant domain. Replacement of D232 with Asn neutralized the negative charge of the carboxylate but preserved the side-chain volume, hydrogen-bonding capacity, and the WT-like loop 3 architecture. The pocket remains accessible to cations, but Asn232 cannot reproduce the negatively charged, multidentate coordination of an Asp carboxylate. Interestingly, the Asn232 side chain adopts a conformation similar to that of Asp232 in Ca^2+^-bound WT C2A, potentially providing a structural explanation for the ability of the D232N mutant to rescue function. D232N therefore preserves the architecture of the site while altering its coordination chemistry. D232A, on the other hand, produces a different structural state. Removal of most of the side chain permits Arg233 to enter the pocket, where it displaces Asp238 and sterically occludes the canonical metal-binding positions. At the same time, the membrane-facing surface is shifted from neutral to positive in the absence of bound metal. Arg233 may partially mimic the electrostatic effect of metal occupancy, although the charge reversal is a cooperative property of the remodeled pocket and does not reproduce Ca^2+^-coordination geometry. The propagation of the D232A rearrangement into residues 189-194 and the β4-β5 hydrophobic core suggests broader intradomain coupling. These structures establish a coupled conformational rearrangement, probably due to allosteric coupling between the Ca^2+^-binding loops and hydrophobic core. A similar observation was reported in octopus SYT1, where a mutation, I248V, on the opposite side of the Ca^2+^-binding sites in the C2A domain, drastically lowered the Ca^2+^-binding affinity^46^. Hence, the idea that Ca^2+^ and membrane interactions with the loops affect only loop structure – without driving conformational changes in the global structure – warrants re-evaluation. Regardless, these structural findings help explain how two distinct neutralizing mutations at position D232 have profoundly different effects on SYT1 function; the D232A mutation traps the protein in a hybrid conformation incompatible with Ca^2+^ binding and membrane penetration, whereas the D232N mutation permits the protein to bind Ca^2+^ even in the absence of Asp. However, these observations are site-specific and do not hold at other positions, as discussed below regarding the C2B domain.

Unsurprisingly, when all five Asp residues in the isolated C2A domain were concurrently mutated to Ala, no Ca^2+^ binding or Ca^2+^-dependent lipid binding activity was observed. Remarkably, in the context of the full-length protein expressed in *syt1* KO neurons, C2A_5A_ rescued ∼70% of evoked synchronous release. Thus, additional Ca^2+^ ligand mutations in C2A can partially overcome the loss-of-function conferred by the single Ala point mutations. The basis for this surprising intragenic suppression is unclear, but several possible explanations exist. First, the C2A_5A_ mutant accumulates on the neuronal plasma membrane, potentially increasing the copy number of this protein at release sites. Since SYT1 oligomerizes, copies on the SV might interact with copies on the plasma membrane, and these interactions might enhance excitation-secretion coupling by this mutant^47^. Second, we emphasize that C2A and C2B functionally interact with one another^25,36^, and these interactions might underlie the ‘dragging model’ presented above. In WT C2A, Ca^2+^ binds to the Asp residues and neutralizes the overall negative charge of the Ca^2+^-and membrane-binding loops, thus favoring electrostatic interactions between positively charged residues in C2A and negatively charged lipids in membranes. In the C2A_5A_ mutant, substituting the five acidic Asp residues with Ala removes the electrostatic repulsion between C2A and anionic phospholipids in the target membrane. This might allow Ca^2+^-bound C2B to pull C2A towards the plasma membrane, consistent with the “dragging” model, thereby facilitating neurotransmitter release (Fig. S10). This model is analogous to the idea presented above regarding the D172A, D232N, and D238A mutants. It can be tested in future experiments by placing a fluorescent reporter in the distal tips of loops 1 and 3 of C2A_5A_ and other mutants, in the complete cytoplasmic domain, and determining whether an adjacent C2B domain drives insertion of the altered C2A domain^45^. Finally, it is possible that C2A_5A_ and other C2A mutants might directly affect inter-domain coupling between C2A and C2B domains, enhancing function in ways we do not yet understand.

Our examination of the putative Ca^2+^ ligands in C2B yielded somewhat different results. Among the five C2B Ca^2+^ ligand mutations, D303A, D363A, and D371A retained some degree of intrinsic Ca^2+^ binding activity via ITC, but only D363A and D371A yielded any detectable degree of residual Ca^2+^-dependent lipid binding activity. In neurons, all C2B CLMs completely disrupted the ability to rescue the extent of evoked synchronous release at physiological [Ca^2+^]_e_, with D371A exhibiting low levels of release at supraphysiological [Ca^2+^]_e_. The inability of D363A to evince glutamate release at high [Ca^2+^]_e_ might be because it has an even lower affinity for Ca^2+^ than D371A (Table S1).

A crucial finding was that every C2B Ca^2+^ ligand mutation, except D371A, completely disrupted the ability of SYT1 to synchronize residual release. This latter finding supports the idea that a key function of C2B is to control the kinetics of exocytosis^40^ and ─ as shown here ─ this function involves the Ca^2+^-sensing activity of this domain. The distinct effects of the D371A mutation are interesting in light of a structural study suggesting that this residue does not serve as a Ca^2+^ ligand ^43^. Our findings suggest that it indeed serves as a ligand but is dispensable for synchronization. Hence, the role of C2B in determining the extent of synchronous release can be partially dissociated from its effects on kinetics. It is tempting to speculate that C2B helps synchronize release by mediating the docking and priming of SVs, thereby enabling them to respond readily to Ca^2+^ ^48^.

For completeness, we concurrently mutated all five acidic ligands in C2B to Ala. In contrast to C2A, the pentuple mutation in C2B abolished the function. This difference provides further evidence that C2A and C2B function somewhat differently. Indeed, C2B is specialized to interact with additional effectors, namely phosphoinositide 4,5-bis-phosphate, t-SNAREs, and – potentially – Ca^2+^-dependent oligomerization^49^.

Our analysis included two disease-associated mutations in C2B, D309A and D309N. This provides a medically relevant comparison between the effects of Ala versus Asn at a Ca^2+^-coordinating residue. While D309A showed no detectable interactions with Ca^2+^ or membranes in biochemical assays, D309N retained partial Ca^2+^- and membrane-binding activity. Structurally, D309A removes the side-chain carboxylate entirely, eliminating a direct Ca^2+^ ligand. In contrast, D309N retains a similarly sized side chain and a carbonyl oxygen that could potentially participate in Ca^2+^ coordination. In the apo structure, however, this oxygen is directed toward an intramolecular interaction with the Asn333 backbone rather than the Ca^2+^-coordinating position observed for Asp309 in the bound WT structures. Interestingly, the Asn309 side chain shows evidence of local conformational heterogeneity, with multiple closely related conformations modeled in the electron density. This local mobility could permit reorientation of the carbonyl oxygen toward a Ca^2+^-competent geometry, providing a structural explanation for the residual Ca^2+^- and membrane-binding activity of D309N. However, both the D309A and D309N mutations completely disrupted SYT1 function in terms of both the extent and synchronization of release, in sharp contrast to the D232A and D232N comparisons in C2A. Although the loss of function in D309A was expected from biochemical studies, the inability of D309N to support release, despite retaining partial Ca^2+^- and membrane-binding activity, suggests that residual binding alone is insufficient. Instead, productive C2B function may require a more ordered Ca^2+^ coordination geometry and membrane-coupled state than D309N can achieve. The complete functional loss of D309N, despite residual biochemical activity, suggests that C2B may be less tolerant than C2A to partial disruption of its Ca^2+^-binding sites, again because C2B engages in more interactions than C2A.

Human patients carrying either D309A or D309N mutation suffer serious developmental challenges. Since these mutations occur in only one allele, we assessed whether they exhibit dominant-negative activity by co-expressing them at equal levels with the WT protein. Both mutants had significant dominant-negative effects on the extent of synchronous release without affecting the kinetics. We attempted to rescue these effects with the drug 4-AP, which inhibits K^+^ channels and enhances Ca^2+^ entry, as described in a previous study^17^. While the extent of synchronous release in the dominant-negative experiments trended upward in the presence of 4-AP, the increase did not reach statistical significance. This lack of effect might be due to the more modest dominant-negative effects described here, with only a ∼30-40% diminution in the peak synchronous glutamate release; this issue will require further study.

In summary, we systematically mutated each putative Ca^2+^ ligand in SYT1 and determined the consequences of these changes using biophysical, structural, and functional approaches. Together, these data establish that all ten proposed residues play crucial roles in coordinating Ca^2+^ ions and triggering exocytosis, overturning dogma that Ca^2+^ binding to C2A is dispensable for function. Indeed, C2A acts as a ‘helper domain’ to enhance the amplitude of evoked synchronous release. We also explored how different Ca^2+^ ligand substitutions can have complex effects on SYT1 structure and function, helping to explain some of the disparities in the literature. Importantly, our analysis revealed that Ca^2+^ binding to C2B underlies the synchronization of release; this warrants further study to address the underlying mechanism(s), which might involve SV docking^48^.

## Materials and Methods

### Constructs used

pLenti-hSynapsin-CRE-WPRE was provided by Fan Wang (Massachusetts Institute of Technology; Addgene ID# 86641)^52^. The glutamate sensor, SF-iGluSnFR.A184V, was provided by Loren Looger (University of California San Diego; Addgene ID# 106175) and was modified as previously described^53^. Rat *syt1* cDNA^6^ was provided by T. C. Sudhof (Stanford University); the D374 mutation was corrected by substitution with a Gly residue. Lentiviral transfer plasmid, FUGW, was provided David Baltimore (California Institute of Technology; Addgene ID# 14883)^54^ and was modified as described previously^53^. SYT1 D232N for crystallization was expressed with an N-terminal GST tag from a modified pGEX-KG vector. All other constructs were cloned into pET-28a(+) with an N-terminal 6xHis tag, and a subset also carried a SUMO or MBP tag downstream of the His tag.

### Recombinant protein expression

Recombinant proteins for biochemical assays were expressed as described previously ^37^. Briefly, all recombinant proteins were expressed in *E. coli* BL21(DE3) cells (NEB, C2527H) at 37 °C until an OD of 0.6 was reached. Expression was induced with 500 μM isopropyl β-D-1-thioga-lactopyranoside (IPTG) (GoldBio, I2481C) followed by overnight growth at 18 °C. Cells were harvested and lysed by sonication, then solubilized in 1% Triton X-100 (Thermo Fisher Scientific, A16046) for 2 h. To minimize proteolytic activity, a protease inhibitor cocktail (PIC) (Roche, 046693132001) was used during the lysis steps. Samples were also incubated with RNase and DNase (10 µg/ml) to prevent nucleic acid-mediated aggregation. Insoluble material was removed by centrifugation at 22,000 x g for 45 min, and the supernatant was incubated with Cobalt TALON affinity resin (Takara, 635653). High-salt washes (0.5-1 M NaCl) were performed to remove bound contaminants. SYT1 C2A or C2B (WT and mutant) domains were eluted from the resin by overnight treatment with 0.5 μM SUMO protease at 4 °C or 200 mM imidazole in 25 mM Tris, pH 7.4, with 500 mM NaCl and 5% glycerol. Purified proteins (>95% purity) were subjected to SDS-PAGE, and protein concentration was determined using bovine serum albumin (BSA) (Jackson ImmunoResearch, 001-000-162) as a standard. Purified proteins were then subjected to dialysis against Chelex 100-treated dialysis buffer (25 mM HEPES pH 7.4, 100 mM KCl).

For crystallization, the cells with the overexpressed SYT1 C2 domain mutant were resuspended in lysis buffer (40 mM HEPES pH 7.4, 150 mM NaCl, 1 mM CaCl_2_), ruptured using a Microfluidics M-110EH microfluidizer, and supplemented with 1 mM PMSF. Lysate was clarified at 45,900 × g (19,500 rpm, Beckman JA-20 rotor) for 1 h at 4°C and incubated overnight (∼12 h) at 4°C on a rocker with pre-equilibrated affinity resin: 20 ml Ni-IDA (His6-Ni Superflow) for His-MBP fusions or 10 ml Glutathione-Sepharose for GST fusions. Ni-IDA columns were washed with 200 ml lysis buffer plus 20 mM imidazole and eluted with 100 ml lysis buffer plus 250 mM imidazole. Eluates were incubated overnight at 4°C with 4 mg TEV protease and 1 mM DTT to release the C2 domain from MBP. GST purifications were performed as described in the previous method ^18^. Briefly, Glutathione-Sepharose columns were washed, protein was eluted with 15 mM reduced glutathione, and cleaved with thrombin for 1 h at RT. Cleaved C2 domains were loaded onto a GE Healthcare SP-Sepharose Fast Flow column on a Bio-Rad NGC system and eluted with a NaCl gradient of Buffer A (lysis buffer) and Buffer B (40 mM HEPES pH 7.4, 1 M NaCl, 1 mM CaCl₂). MBP and GST did not bind the resin, and TEV protease eluted before the C2 domain. Fractions were selected by SDS-PAGE (Bio-Rad Any-kD Mini-PROTEAN TGX Stain-Free gels), pooled, concentrated to 1 ml (Amicon Ultra-15, 10 kDa MWCO; UFC901024), and further purified by size exclusion chromatography on a Superdex-75 column in 40 mM HEPES pH 7.4, 300 mM NaCl, 1 mM CaCl₂. Purity was confirmed by SDS-PAGE, and concentration was determined by A₂₈₀ using extinction coefficients calculated with ExPASy ProtParam. Purified protein was flash-frozen in liquid nitrogen and stored at −80 °C.

### Isothermal titration calorimetry (ITC)

ITC measurements were performed using a MicroCal iTC200 calorimeter (Malvern Panalytical, UK), following the procedures described previously^37^. SYT1 C2A or C2B domains (50 µM) were titrated with 5 mM Ca^2+^. All solutions were prepared in Chelex 100-treated dialysis buffer (25 mM HEPES pH 7.4, 100 mM KCl) and degassed at 25 °C prior to each experiment. Each titration consisted of an initial 2 µl injection of Ca^2+^ followed by 18 successive 4 µl injections, with stirring at 750 rpm at 25 °C. Heats of dilution were corrected by subtracting a blank titration of Ca^2+^ into buffer alone. Titrations were performed in triplicate, and binding constants were obtained by fitting the data to a two-or three-site binding model using MicroCal Origin 2020 software (Malvern Panalytical, UK).

### Co-sedimentation assay

Co-sedimentation assays were performed as previously described^36^. Briefly, isolated WT or mutant SYT1 C2A and C2B domains (2.5 µM) were mixed with liposomes (50 μl of 1 mM total lipids; composition: 20% DOPS, 80% DOPC; 100 nm diameter) in HEPES buffer (25 mM HEPES pH 7.4, 100 mM KCl) at the indicated free [Ca^2+^]. Samples were centrifuged at 170,000 × g for 60 min at 4 °C in an Optima MAX-E tabletop ultracentrifuge (Beckman Coulter). The supernatant from each sample was collected and analyzed by SDS-PAGE with ReadyBlue staining (Sigma-Aldrich). Unbound protein was quantified by densitometry, and the percentage of bound protein was calculated from the depletion of protein in the supernatant. Each experiment was independently repeated at least three times, using at least two protein preparations and three liposome preparations

### Cell culture

HEK293T cells (ATCC, CRL-11268) for lentivirus production were cultured in Dulbecco’s Modified Eagle Medium (DMEM) with high glucose (Gibco, 11965092), supplemented with 10% fetal bovine serum (FBS; R&D Systems, S11550H) and penicillin-streptomycin (Thermo Fisher Scientific, MT-30-001 CI), and passaged upon 80% confluency.

Mouse hippocampal neurons from the *syt1*-floxed mouse strain^55^ were isolated at postnatal day 0 or 1. Hippocampi were dissected and maintained in chilled Hibernate A media (BrainBits; HA) supplemented with N-21 max (2%, R&D Systems, AR008) and Glutamax (2 mM Gibco; 35050061). Neuronal tissue was then incubated with 0.25% trypsin (Corning; 25–053 CI) for 25 min at 37 °C, washed 5 times with DMEM (Gibco, 11965092) supplemented with 10% FBS plus penicillin-streptomycin, and triturated until homogeneous. Neurons were counted using a Scepter 3.0 automated cell counter (Millipore Sigma; PHCC340) and plated in DMEM supplemented with 10% FBS and pen/strep at a density of 200K cells per 18 mm coverslip; the coverslips were pre-coated with poly-D-lysine and EHS laminin (Thermo Fisher Scientific; 23017015) for 2 hours prior to plating. Neurons were incubated at 37 °C, 5% CO_2._ DMEM was replaced with Neurobasal-A (Thermo Fisher; 10888–022), supplemented with N-21 max, Glutamax and pen/strep after 1 hour. Cultures were supplemented with fresh Neurobasal-A medium on day in vitro (DIV) 5 and DIV11 and were used for experiments between DIV14-16.

### Lentivirus prep and transduction

Lentivirus production was performed using the protocol previously described^53,56^. Neurons were transduced with pLenti-hSynapsin-CRE-WPRE at DIV1 to express Cre recombinase. Lentiviruses expressing iGluSnFR.A184V and SYT1 mutants were added on DIV5. iGluSnFR lentivirus was titrated based on fluorescence. All other lentiviruses were titrated via immunoblot analysis for each production batch.

### Immunoblotting

Cultures were washed with ice-cold DPBS to remove interfering phenols and albumins, then lysed in 150 μl lysis buffer (2% SDS, 1% Triton X-100, and 10 mM EDTA in PBS, supplemented with protease inhibitors: 2 mM PMSF, 2 μg/ml aprotinin, 2 μg/ml pepstatin A, and 2 μg/ml leupeptin). Lysates were centrifuged at 20,000 × g for 10 min at 4°C, and protein concentration in the supernatant was determined using bicinchoninic acid (BCA) assay (Thermo Fisher Scientific, 23227). The remaining supernatant was mixed with 4× Laemmli sample buffer (Bio-Rad, 1610747) containing 10% β-mercaptoethanol (Bio-Rad, 1610710XTU) and heated at 75 °C for 5 min. Samples containing 5 μg of total protein were resolved by SDS-PAGE on 12% TGX Stain-Free gels (Bio-Rad, 5678043). Following a 5 min activation at 300 nm, gels were imaged on a ChemiDoc MP system (Bio-Rad), and the resulting total protein signal was used as a loading control. Proteins were transferred to PVDF membranes (EMD Millipore, IPFL00010) using the Trans-Blot Turbo Transfer System (Bio-Rad) at constant current (240 mA). Membranes were blocked for 1 h in 5% nonfat milk in Tris-buffered saline with 0.1% Tween-20 (TBST) and incubated overnight at 4 °C with anti-SYT1 antibody (1:1000; Synaptic Systems, 105 011) in 2% nonfat milk in TBST. After three washes in TBST, membranes were incubated for 1 h at room temperature with HRP-conjugated anti-mouse secondary antibody (Bio-Rad 1706516; dilution 1:10000), followed by three additional TBST washes. Immunoblots were developed with Luminata Forte Western HRP Substrate (EMD Millipore, ELLUF0100) and imaged on a ChemiDoc MP system (Bio-Rad). Band intensities were quantified by densitometry, and contrast was linearly adjusted for presentation in Fiji^57^.

### Immunocytochemistry

Samples were prepared as previously described^58^. Briefly, dissociated cultures were fixed with ice-cold methanol, permeabilized with 0.2% saponin, blocked (0.04% saponin, 10% goat serum, and 1% BSA in PBS), and then immunostained with mouse anti-SYT1 (1:500; Synaptic Systems, 105 011) and guinea pig anti-SYP (1:500; Synaptic Systems, 101 004) antibodies in 0.1% BSA and 0.04% saponin in PBS overnight at 4 °C. The following morning, coverslips were rinsed three times with PBS and incubated with secondary antibodies (goat anti-mouse IgG2a AF 488, Thermo Scientific A-21131, 1:500; goat anti-guinea pig AF 647, Thermo Scientific A-21450, 1:500) diluted in PBS containing 0.1% BSA and 0.04% saponin for 1 hr. Then, coverslips were rinsed three times with PBS and mounted on microscope slides (Thermo Fisher Scientific, 22-178277) using ProLong Glass Antifade with Mountant with NucBlue Stain (Thermo Fisher Scientific, P36981). Coverslips were imaged using a Zeiss 880 Airyscan LSM microscope with a 63X/1.4 NA oil objective at room temperature, and images were processed with automatic Airyscan deconvolution settings. Co-localization analysis was performed using Fiji for ImageJ and Just Another Colocalization Plugin (JaCoP)^57,59^.

### iGluSnFR imaging and analysis

Glutamate imaging and quantification were performed as previously described using SF-iGluSnFR.A184V with minor modifications^53^. Briefly, experiments were performed using an Olympus IX83 inverted microscope equipped with an X-Cite 120 LED (Lumen Dynamics) and an ORCA-Fusion CMOS camera (Hamamatsu Photonics). Field stimulation was used to trigger a single action potential; 150 frames (10 ms exposure time) were collected over 1.5 s with 2×2 binning. Extracellular imaging media (140 mM NaCl, 5 mM KCl, 1-10 mM CaCl_2_, 1 mM MgCl_2_, 5.5 mM glucose, 20 HEPES, pH 7.4, in mM), with an osmolarity of 300-320 mOsm, was supplemented with 50 μM D-AP5 (Abcam, ab120003), 20 μM CNQX (Abcam; ab120044), and 100 μM Picrotoxin (Tocris; 1128) to block recurrent activity. All experiments were carried out at 33-34 °C, with neurons at DIV14-16. Temperature and humidity were controlled by a Tokai incubation controller and chamber (Tokai Hit, PPZI3).

A custom ImageJ plugin was used to identify iGluSnFR regions of interest (ROIs)^53^. Intensity values obtained from the plugin were imported to AxographX1.8.0 (Axograph Scientific), where traces were normalized to the initial 500 ms pre-stimulus baseline and corrected with a two-tailed background subtraction. Glutamate peaks were defined as fluorescence values ≥ 4x the standard deviation (SD) of the pre-stimulus baseline signals.

### Crystallization, data collection, and processing

Purified SYT1 C2A D232A was initially crystallized at 25 mg/ml using the sitting drop vapor diffusion method with 200 nl protein and 200 nl well solution, left at 20 °C. Crystallization experiments for SYT1 C2A D232A were set up using the TTP LabTech Mosquito in the UW-Madison Crystallography Core facility. These crystals, grown in 0.25 M ammonium sulfate, 28% PEG 8000, 0.1 M sodium acetate, pH 4.8, were resuspended in 100 µL of well solution and crushed using a Crystal Crusher (Hampton Research: HR4-216). The seed stock was diluted 1:5 and used to obtain the final protein crystals, again using the sitting drop vapor diffusion method with 200 nl protein at 10 mg/mL and 200 nl well solution (0.25 M ammonium sulfate, 28% PEG 8000, 0.1 M sodium acetate pH 4.8), and 50 nl of the 1:5 diluted crystal seeds. These crystals were cryoprotected with 20% glycerol, picked up with nylon loops, and quickly frozen in liquid N_2_. Next, these crystals were shipped to the Advanced Photon Lightsource (APS) for data collection at beamline 23-ID-D at 90 K with a wavelength of 1.0332 Å. Final X-ray diffraction data were processed using the auto-processing feature of APS with the xia2 and DIALS pipeline^60^.

Purified C2A D232N was crystallized at 25 mg/ml using the hanging drop vapor diffusion method with 2 µl protein and 2 µl well solution. The crystals were grown at 17 °C in 2.3 M lithium sulfate and 100 mM HEPES pH 7.0. These crystals were picked using nylon loops, and subsequently frozen with liquid N_2_ and shipped to Stanford Synchrotron Radiation Lightsource (SSRL). The final dataset was collected at beamline 9-1 at 90 K with a wavelength of 0.818 Å. Final X-ray diffraction data were processed using the HKL3000 software package^61^.

Purified SYT1 D309A was crystallized at 22 mg/ml using the hanging drop vapor diffusion method with 2 µl protein and 2 µl well solution. The crystals were grown at 20 °C with 0.07 M ammonium sulfate, 25% PEG 4000, and 15% glycerol, and cryoprotected with 0.1 M ammonium sulfate, 25% PEG 4000, and 30% glycerol. Purified SYT1 D309N was crystallized at 21 mg/ml using the hanging drop vapor diffusion method with 2 µl protein and 4 µl well solution. The crystals were grown at 20 °C in a solution containing 0.07 M ammonium sulfate, 25% PEG 4000, and 15% glycerol, and cryoprotected with Perfluoropolyether Cryo Oil (Hampton Research: HR2-814). The crystals were picked using nylon loops and subsequently frozen with liquid N_2_. All C2B mutant crystals were shipped to Stanford Synchrotron Radiation Lightsource (SSRL), and datasets were collected at beamline 9-2 at 90 K with a wavelength of 0.9795 Å. Final X-ray diffraction data were processed using the autoxds script from SSRL. Briefly, XSCALE (part of the XDS package) scales the data, POINTLESS determines the correct space group, AIMLESS scales and merges the reflections, and finally TRUNCATE is used to convert the intensities to amplitudes^62^.

All X-ray crystal structures were solved by molecular replacement with PHENIX via the Phaser-MR GUI^63,64^ using structural model 6TZ3 for the C2B domains, D309A and D309N, and 4WEE for the C2A domains, D232A and D232N. All refinement was done using the PHENIX software package and the PDB-REDO web server^64,65^ while model building was performed using COOT ^66^. Final structural models were deposited into the Worldwide Protein Data Bank (wwPDB) and received the following PDB IDs: 9ZH6 (D232A), 9ZKH (D232N), 9ZJL (309A), 9ZF1 (D309N).

### Detailed clinical histories for patients D309A and D309N

The first proband, a now 7-year-old female, was born at term to a P1G1 mother via emergency C-section. Parents were non-consanguineous, with a history of autism and ADHD on the maternal side. Laryngomalacia was first noted at one month of age and has since resolved. At six months of age, parents observed a delay in reaching developmental milestones, and the first seizure was reported at one year of age, following a febrile incident. Rapid brain MRI was unremarkable except for positional plagiocephaly. The patient currently exhibits global developmental delays, with developmental age corresponding to a 4-month-old. She can roll side to side, sit with assistance, grab objects, and feed herself from a bottle. She is non-verbal with some vocalizations. Neurological exam shows the patient to be normocephalic with dysconjugate gaze, hypermobile joints, diffuse hypotonia, involuntary hyperkinetic movements with no EEG correlation, and dysphagia. At 1.5 years of age, whole-exome sequencing (GeneDx XomeDx) revealed a heterozygous missense mutation in the *syt1* gene, pD310A (corresponding to D309A in the mouse), which was not present in the proband’s mother. EEG demonstrated abnormal brain activity but did not correlate with seizures. The patient was put on anti-seizure medications in the past but has been off all seizure medications for the past 3 years without any concerning seizure activity.

The second proband, a now 13-year-old male, was born to non-consanguineous parents. At 2 years of age, the patient presented with ataxia, delayed speech and language development, developmental regression, focal-onset seizures, hyperlordosis, hypotonia, motor delay, scoliosis, and skeletal muscle atrophy. Current symptoms include difficulty in speaking, epileptic anomaly, and slow motor development. Whole exome sequencing (CENTOGENE, CentoXome® Trio) identified a *de novo* heterozygous missense mutation in the *syt1* gene, pD310N (corresponding to D309N in the mouse), which was examined in the present study. In addition, a heterozygous mutation in CYP24A1 (E143del) was also found. CYP24A1 encodes the enzyme 24-hydroxylase, which is involved in vitamin D catabolism and Ca^2+^ homeostasis. E143del in CYP24A1 leads to complete loss of function and is associated with idiopathic infantile hypercalcemia^67^. Symptoms include profound hypercalcemia, suppressed intact parathyroid hormone, hypercalciuria, and medullary nephrocalcinosis. The mutation is suspected to be inherited in an autosomal recessive manner. Given the patient’s heterozygosity, we suspect this mutation has minimal impact on the patient’s symptoms. However, a definite negation of the causal effect of this mutation is outside the scope of the current study.

## Acknowledgments

We are indebted to Dr. Latisha McLaurin, Texas Tech University Health Sciences Center, Lubbock, and the School of Medicine and Public Health Informatics and Information Technology, UW-Madison, for their help in obtaining and de-identifying patient information. We also thank all members of the Chapman lab, as well as former members M. Seibert and E. Watson, for valuable discussions and feedback on this manuscript. For SYT1 C2A D232N (9ZKH), SYT1 C2B D309A (9ZJL), and SYT1 C2B D309N (9ZF1), data were collected at SSRL BL 11-1, 9-2, and 9-2, respectively. For SYT1 C2A D232A (9ZH6) crystallization, data collection was conducted by the Department of Biochemistry Collaborative Crystallography Core at the University of Wisconsin, which is supported by user fees and the department. Data were collected at APS 23-ID-D. ERC is an Investigator of the Howard Hughes Medical Institute (HHMI).

## Funding

National Institutes of Health grant R01MH061876, R35NS136306 (ERC) Howard Hughes Medical Institute (ERC)

## Author contributions

Conceptualization: SM, NM, MJD, RBS, ERC Data curation: SM

Formal analysis: SM, NM, MJD, RBS, ERB Investigation: SM, NM, MJD, ERB, ERC Methodology: SM, NM, MJD, MW, ERB, AV, ERC

Software: SM, NM, MJD Supervision: SM, ERC Visualization: SM

Writing – original draft: SM, ERC

Competing interests: The authors declare no competing interests.

## Data and materials availability

All gels, blots, and statistical information are available in the source data file. Raw data will be made available in a publicly available repository Plasmids are made available on Addgene. All other information is available in the main text or supplementary material. X-ray crystallography structures have been deposited at the RCSB PDB and are publicly available as of the date of publication. De-identified patient information can be requested from the corresponding author, subject to approval by the University of Wisconsin Office of Compliance.

## Ethics Statement

Animal care and use in this study were conducted under guidelines set by the NIH’s Guide for the Care and Use of Laboratory Animals handbook. Protocols were reviewed and approved by the Animal Care and Use Committee at UW-Madison (Laboratory Animal Welfare Public Health Service Assurance Number: A3368-01). This study was conducted in accordance with the Declaration of Helsinki and applicable U.S. regulations. The work reports only patient history: no experimental interventions, procedures, or additional data collection were performed for this study. Legal guardians of the patients provided electronic informed consent for the research use and publication of their clinical histories, under the oversight of the University of Wisconsin Office of Compliance. Patient history information was obtained from clinical records provided by the family. The information reported here was de-identified by the research team using the HIPAA Safe Harbor method, with the removal of all 18 specified identifiers, and de-identification was independently verified by the institution’s honest broker office.

## Notes

http://datadryad.org/share/LINK_NOT_FOR_PUBLICATION/S_x3q3uWwa6dgfuTbJ7m9QZ153DS69ObFF6eQFEWR6o

### Competing Interest Statement

The authors have declared no competing interest.

