## Supplementary Information for "Functional mapping of synaptotagmin 1 Ca^2+^ ligands that mediate excitation-secretion coupling in neurons"

**Tables S1 to S3**

**Figs. S1 to S20**

**References**

**SYT1 C2A**

| Dissociation constant | WT | D232N |
|---|---|---|
| # binding sites | 3 | 2 |
| $K_{d1}$ (µM) | 68.2 ± 4.20 | 231 ± 18.8 |
| $K_{d2}$ (µM) | 103 ± 34.0 | 1660 ± 125 |
| $K_{d3}$ (µM) | 1130 ± 120 | |

**SYT1 C2B**

| Dissociation constant | WT | D303A | D363A | D371A | D309N |
|---|---|---|---|---|---|
| # binding sites | 2 | 2 | 2 | 2 | 2 |
| $K_{d1}$ (µM) | 180 ± 4.90 | 385 ± 11.1 | 435 ± 12.8 | 298 ± 14.0 | 359 ± 32.6 |
| $K_{d2}$ (µM) | 240 ± 16.0 | 510 ± 89.3 | 754 ± 64.0 | 473 ± 91.2 | 618 ± 87.6 |

| | ΔH (cal/mol) | ΔS (cal/mol/K) |
|---|---|---|
| **C2A WT** | 577 ± 194 | 21.0 ± 0.75 |
| | 309 ± 108 | 19.4 ± 0.60 |
| | 361 ± 91 | 14.7 ± 0.35 |
| **D232N** | 395 ± 117 | 18.0 ± 0.35 |
| | 381 ± 343 | 14.0 ± 1.3 |
| **C2B WT** | -427 ± 108 | 15.7 ± 0.4 |
| | -585 ± 300 | 14.6 ± 1.14 |
| **D303A** | -305 ± 114 | 14.6 ± 0.4 |
| | -641 ± 14 | 12.9 ± 0.32 |
| **D363A** | -411 ± 140 | 14.0 ± 0.46 |
| | -643 ± 59 | 12.1 ± 0.35 |
| **D371A** | -268 ± 124 | 15.2 ± 0.4 |
| | -428 ± 109 | 13.8 ± 0.6 |
| **D309N** | -427 ± 63 | 14.3 ± 0.38 |
| | -714 ± 199 | 12.3 ± 0.4 |

**Table S1:** Dissociation constants ($K_d$ values) and thermodynamic properties (H, enthalpy; S, entropy) of $Ca^{2+}$ binding at 25 °C to the indicated SYT1 constructs measured using ITC. Data are mean values ± SD; N = 3.

| | SYT1 C2A D232A | SYT1 C2A D232N | SYT1 C2B D309A | SYT1 C2B D309N |
|---|---|---|---|---|
| **PDB ID** | 9ZH6 | 9ZKH | 9ZJL | 9ZF1 |
| **Wavelength (Å)** | 1.033 | 0.818 | 0.9795 | 0.9795 |
| **Resolution range (Å)** | 59.17 - 2.74 (2.84 - 2.74) | 22.68 - 1.2 (1.24 - 1.2) | 27.85 - 1.07 (1.11- 1.07) | 35.19 - 1.05 (1.09 - 1.05) |
| **Space group** | P 61 2 2 | P 1 21 1 | P 32 2 1 | P 32 2 1 |
| **Unit cell** | 68.3 68.3 268.5 90 90 120 | 41.7 38.5 44.1 90 98.31 90 | 54.6 54.6 103.4 90 90 120 | 54.8 54.8 105.0 90 90 120 |
| **Total reflections** | 111097 (11307) | 315988 (25855) | 2852975 (165161) | 2770412 (93886) |
| **Unique reflections** | 10546 (1014) | 43200 (4037) | 79112 (7132) | 85545 (7486) |
| **Multiplicity** | 10.5 (11.2) | 7.3 (6.4) | 36.1 (21.7) | 32.4 (11.3) |
| **Completeness (%)** | 99.7 (98.4) | 99.2 (93.8) | 99.1 (91.7) | 98.8 (88.5) |
| **Mean I/sigma(I)** | 2.77 (0.55) | 26.16 (6.22) | 26.02 (0.40) | 26.47 (0.43) |
| **Wilson B-factor** | 43.66 | 10.44 | 12.92 | 13.29 |
| **R-merge** | 0.62 (3.50) | 0.036 (0.28) | 0.25 (3.40) | 0.78 (2.37) |
| **R-meas** | 0.65 (3.67) | 0.039 (0.30) | 0.25 (3.48) | 0.79 (2.49) |
| **R-pim** | 0.2 (1.10) | 0.014 (0.12) | 0.040 (0.72) | 0.13 (0.72) |
| **CC1/2** | 0.94 (0.29) | 1 (0.96) | 0.99 (0.41) | 0.7 (0.42) |
| **CC*** | 0.99 (0.67) | 1 (0.99) | 0.99 (0.76) | 0.91 (0.80) |
| **Reflections used in refinement** | 10511 (998) | 43189 (4037) | 78608 (7130) | 84726 (7480) |
| **Reflections used for R-free** | 500 (39) | 1996 (184) | 1988 (177) | 1998 (183) |
| **R-work** | 0.2730 (0.4220) | 0.1428 (0.1488) | 0.1531 (0.3336) | 0.1680 (0.3757) |
| **R-free** | 0.3047 (0.3909) | 0.1622 (0.1859) | 0.1692 (0.3739) | 0.1922 (0.3779) |
| **CC (work)** | 0.907 (0.465) | 0.967 (0.964) | 0.919 (0.731) | 0.815 (0.658) |

| | | | | |
|---|---|---|---|---|
| **CC (free)** | 0.827 (0.470) | 0.978 (0.940) | 0.956 (0.648) | 0.884 (0.703) |
| **Number of non-hydrogen atoms** | 2085 | 1456 | 1555 | 1653 |
| **macromolecules** | 2064 | 1245 | 1359 | 1445 |
| **ligands** | 21 | 18 | 36 | 37 |
| **solvent** | 0 | 193 | 168 | 181 |
| **Protein residues** | 254 | 136 | 149 | 149 |
| **RMS (bonds)** | 0.004 | 0.018 | 0.011 | 0.018 |
| **RMS (angles)** | 0.77 | 1.67 | 1.14 | 1.49 |
| **Ramachandran favored (%)** | 95.20 | 97.01 | 97.96 | 96.60 |
| **Ramachandran allowed (%)** | 4.00 | 2.24 | 2.04 | 2.72 |
| **Ramachandran outliers (%)** | 0.80 | 0.75 | 0.00 | 0.68 |
| **Rotamer outliers (%)** | 3.10 | 0.73 | 2.60 | 1.23 |
| **Clashscore** | 8.47 | 7.56 | 4.61 | 4.69 |
| **Average B-factor** | 42.48 | 16.58 | 21.70 | 22.02 |
| **macromolecules** | 42.28 | 14.69 | 19.49 | 20.54 |
| **ligands** | 61.55 | 26.42 | 52.35 | 28.57 |
| **solvent** | | 27.89 | 34.47 | 32.86 |

**Table S2:** Data collection and refinement statistics. Statistics for the highest-resolution shell are shown in parentheses.

| PDB | $Ca^{2+}$ numbering in PDB file | Coordination mode | D309 oxygen(s) | Distance(s), Å |
|---|---|---|---|---|
| 1TJX | 1420 | monodentate | OD1 | 2.50 |
| 1UOV | 1420 | monodentate | OD2 | 2.35 |
| 1UOV | 1421 | bidentate | OD1, OD2 | 2.37, 2.60 |
| 7TX9 | 501 | monodentate | OD2 | 2.59 |

**Table S3.** D309 coordination in $Ca^{2+}$-bound reference structures.

| | Loop 1 | | | | | | | | | | | | Loop 3 | | | | | | | | |
|---|---|---|---|---|---|---|---|---|---|---|---|---|---|---|---|---|---|---|---|---|---|
| **SYT1 C2A** | 167 | 168 | 169 | 170 | 171 | 172 | 173 | 174 | 175 | 176 | 177 | 178 | 230 | 231 | 232 | 233 | 234 | 235 | 236 | 237 | 238 |
| Rat | E | L | P | A | L | D | M | G | G | T | S | D | D | F | D | R | F | S | K | H | D |
| Mouse | E | L | P | A | L | D | M | G | G | T | S | D | D | F | D | R | F | S | K | H | D |
| Human | E | L | P | A | L | D | M | G | G | T | S | D | D | F | D | R | F | S | K | H | D |
| Chicken | E | L | P | A | L | D | M | G | G | T | S | D | D | F | D | R | F | S | K | H | D |
| Bovine | E | L | P | A | L | D | M | G | G | T | S | D | D | F | D | R | F | S | K | H | D |
| Drosophila | E | L | P | A | L | D | M | G | G | T | S | D | D | F | D | R | F | S | K | H | D |
| *C. elegans* | D | L | P | G | M | D | M | S | G | T | S | D | D | F | D | R | F | S | K | H | D |

| | Loop 1 | | | | | | | | | | | | | | Loop 3 | | | | | | | | |
|---|---|---|---|---|---|---|---|---|---|---|---|---|---|---|---|---|---|---|---|---|---|---|---|
| **SYT1 C2B** | 297 | 298 | 299 | 300 | 301 | 302 | 303 | 304 | 305 | 306 | 307 | 308 | 309 | 310 | 363 | 364 | 365 | 366 | 367 | 368 | 369 | 370 | 371 |
| Rat | K | N | L | K | K | M | D | V | G | G | L | S | D | P | D | Y | D | K | I | G | K | N | D |
| Mouse | K | N | L | K | K | M | D | V | G | G | L | S | D | P | D | Y | D | K | I | G | K | N | D |
| Human | K | N | L | K | K | M | D | V | G | G | L | S | D | P | D | Y | D | K | I | G | K | N | D |
| Chicken | K | N | L | K | K | M | D | V | G | G | L | S | D | P | D | Y | D | K | I | G | K | N | D |
| Bovine | K | N | L | K | K | M | D | V | G | G | L | S | D | P | D | Y | D | K | I | G | K | N | D |
| Drosophila | K | N | L | K | K | M | D | V | G | G | L | S | D | P | D | Y | D | R | I | G | T | S | E |
| *C. elegans* | K | N | L | K | K | M | D | V | G | G | L | S | D | P | D | Y | D | K | L | G | S | N | D |

**Fig. S1: Conservation of Asp residues in loops 1 and 3 in SYT1.** Alignment of putative $Ca^{2+}$ ligands in SYT1 from multiple species. Residues here are numbered using SYT1 from *Rattus norvegicus*. All ten Asp residues are highly conserved across species, except in *Drosophila,* where the last Asp residue in C2B is substituted with another acidic residue, Glu. Aligned using Clustal Omega[1,2].

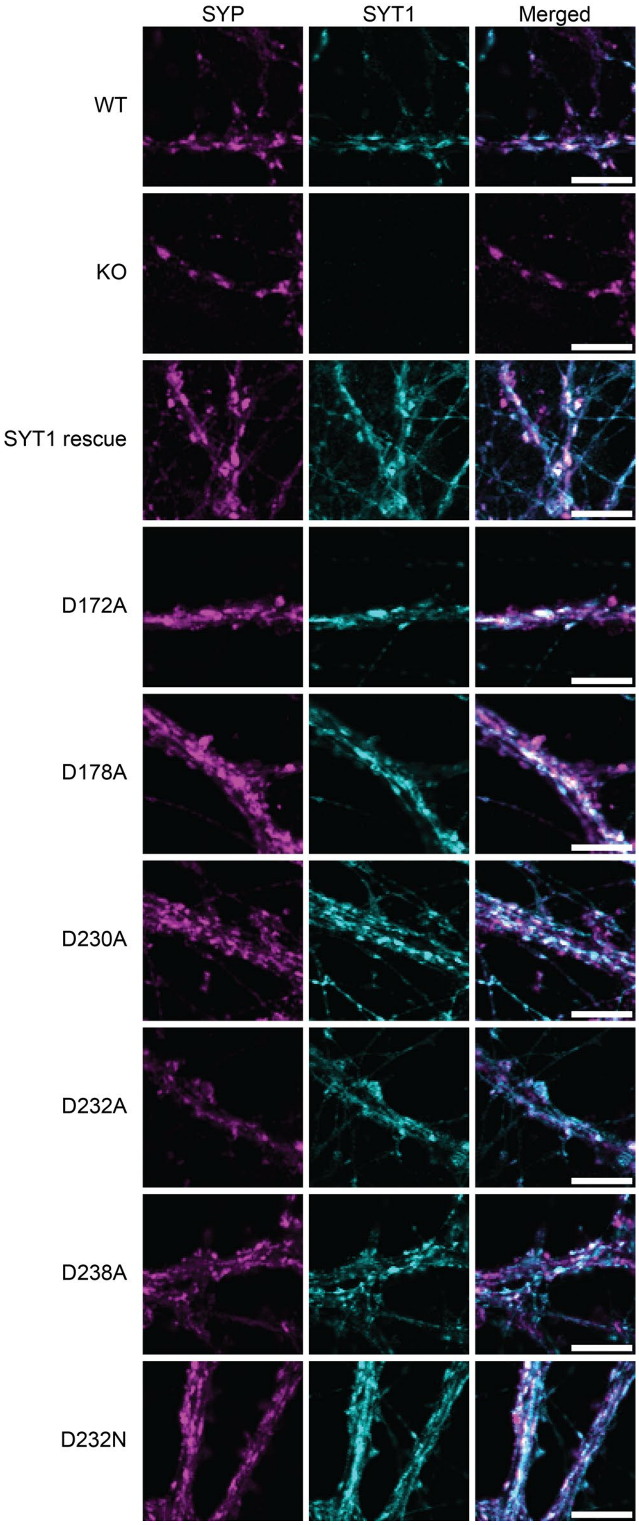


**Fig. S2: C2A $Ca^{2+}$ ligand mutants are localized to SVs.** Representative images showing co-localization of C2A CLMs with the SV protein synaptophysin (SYP). Scale bar, 5 µm.

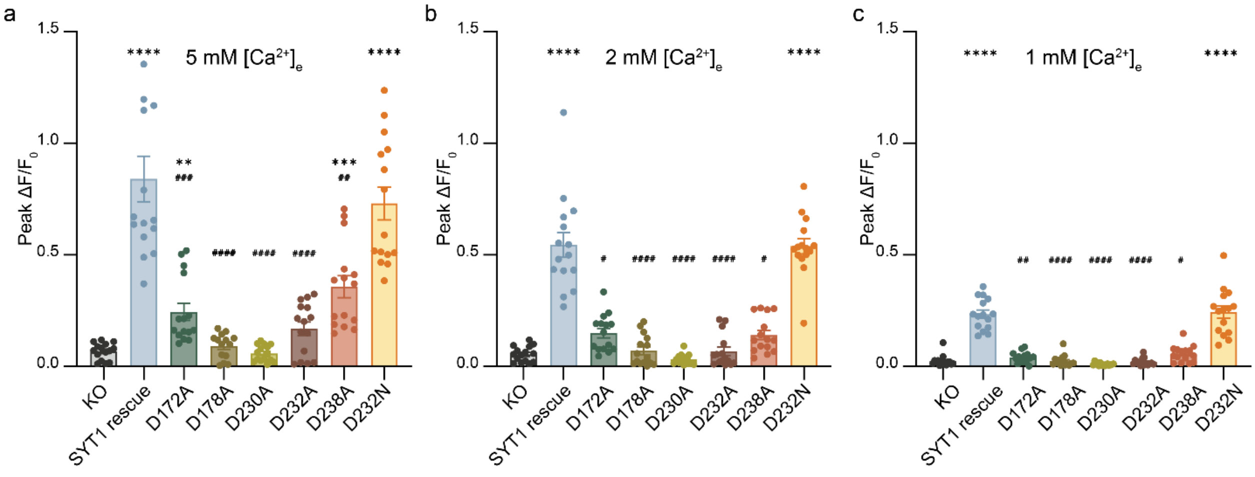


**Fig. S3: C2A CLMs disrupt glutamate release at lower $[Ca^{2+}]_e$:** Peak iGluSnFR $\Delta F/F_0$ of C2A CLMs at 5, 2 and 1 μM $[Ca^{2+}]_e$. Each data point represents a single FOV, n = 15 FOVs, N = 3 independent trials; error bars indicate SEM. Statistical significance was determined using the Brown-Forsythe ANOVA test, with Dunnett's T3 multiple comparisons test. For all panels, ****, #### $p < 0.0001$; ***, ### $p < 0.001$; **, ## $p < 0.01$; *,# $p < 0.05$.

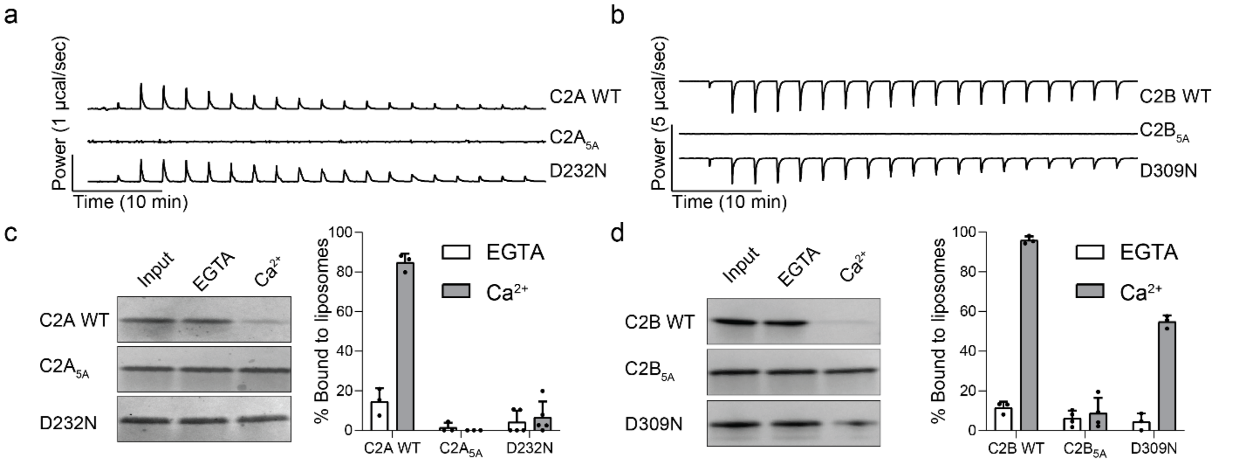


**Fig. S4: a,** Representative isothermal titration calorimetry (ITC) traces showing the heat of $Ca^{2+}$ binding to isolated WT C2A, $C2A_{5A}$, and D232N, **b,** Same as panel (a), but for WT C2B, $C2B_{5A}$ and D309N. Dissociation constants are reported in Supplementary Table 1. **c,** Left: Representative SDS-PAGE gels showing the supernatant from WT C2A, $C2A_{5A}$, and D232N, C2A-domain•liposome co-sedimentation experiments, in the absence (EGTA) and presence of $Ca^{2+}$. Input refers to protein-only control. Right: Quantification of %protein bound to liposomes, in the absence and presence of $Ca^{2+}$. N ≥ 3. **d,** Same as panel (c), but for WT C2B, $C2B_{5A}$ and D309N. The Mann-Whitney test was performed to assess rank separation.

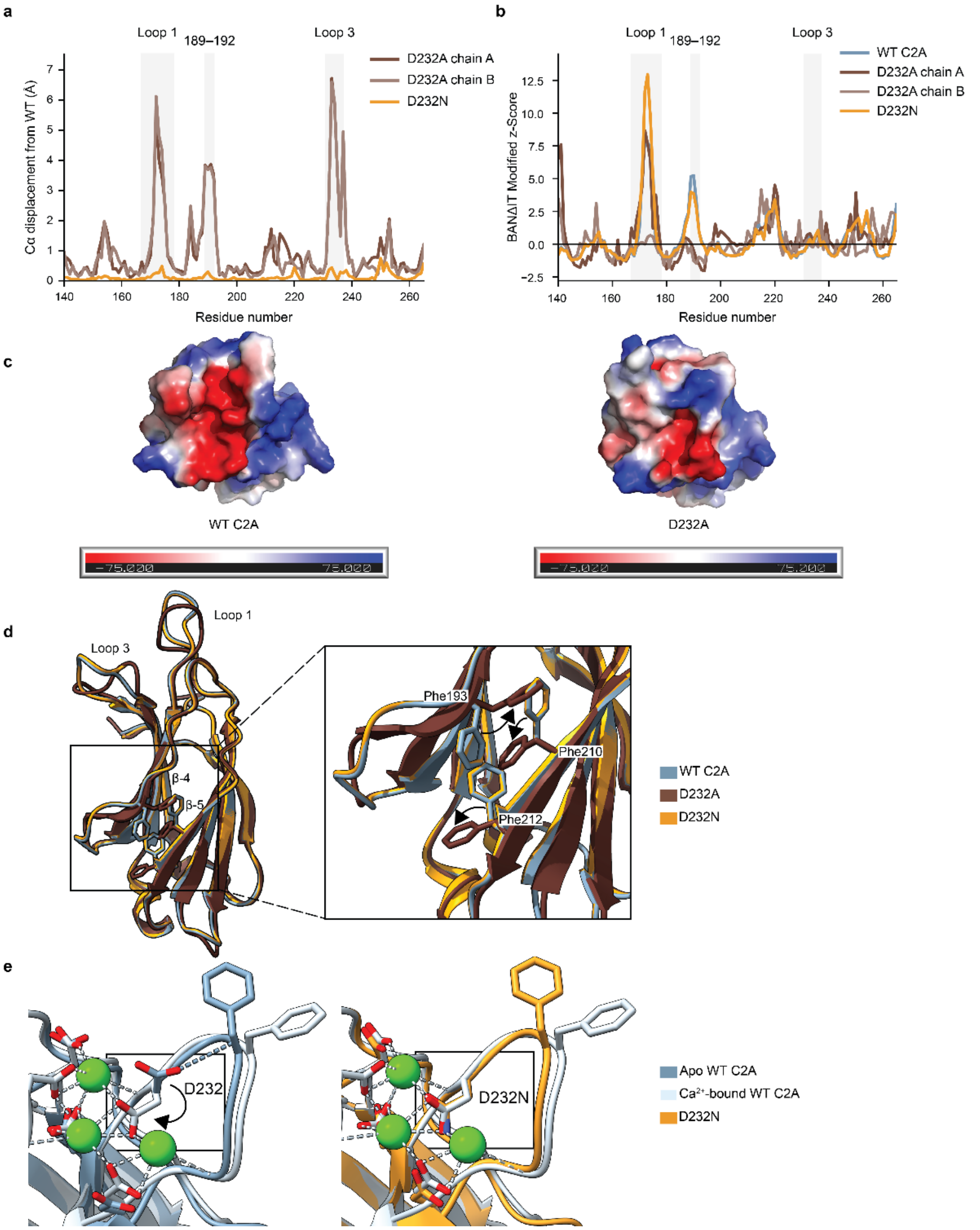


**Fig. S5: Loop 1 displacement and normalized mobility in WT, D232A, and D232N.** **a,** Per-residue Cα displacement of D232A and D232N relative to WT SYT1 C2A following alignment of the conserved β-sandwich core. The largest modeled differences in D232A are localized to loops 1 (E167-D178) and 3 (D230-D238), whereas D232N remains unchanged compared to apo WT. **b,** BANΔIT Modified z-Score profiles calculated from Cα B-factors for WT, D232A, and D232N. Shaded regions indicate loop 1 and loop 3. Loop 1 is intrinsically mobile in WT, D232N, and D232A chain A, while the same region in D232A chain B is comparatively well ordered. **c,** Electrostatic surface potential of WT C2A and D322A showing the change in the membrane-facing surface. Red, blue, and white in the color ramp depict negative, positive, and neutral charges, respectively. The electrostatic surface was rendered using PyMOL[3]. **d,** Superimposition of the asymmetric units of SYT1 C2A D232A (PDB: 9ZH6, chain B) and SYT1 C2A D232N (PDB: 9ZKH), compared with WT SYT1 C2A (PDB:4WEE) showing a flip in the adjacent hydrophobic core. Magnified view depicting altered orientations of Phe193, Phe210, and Phe212. **e,** Superimposition of the asymmetric units of comparing SYT1 C2A D232N (PDB: 9ZKH) to apo WT SYT1 C2A (PDB:4WEE) (left) and $Ca^{2+}$-bound WT SYT1 C2A (PDB:1BYN) (right), highlighting the rotation of Asn232. Normalized Cα B-factors were computed using BANΔIT[4].

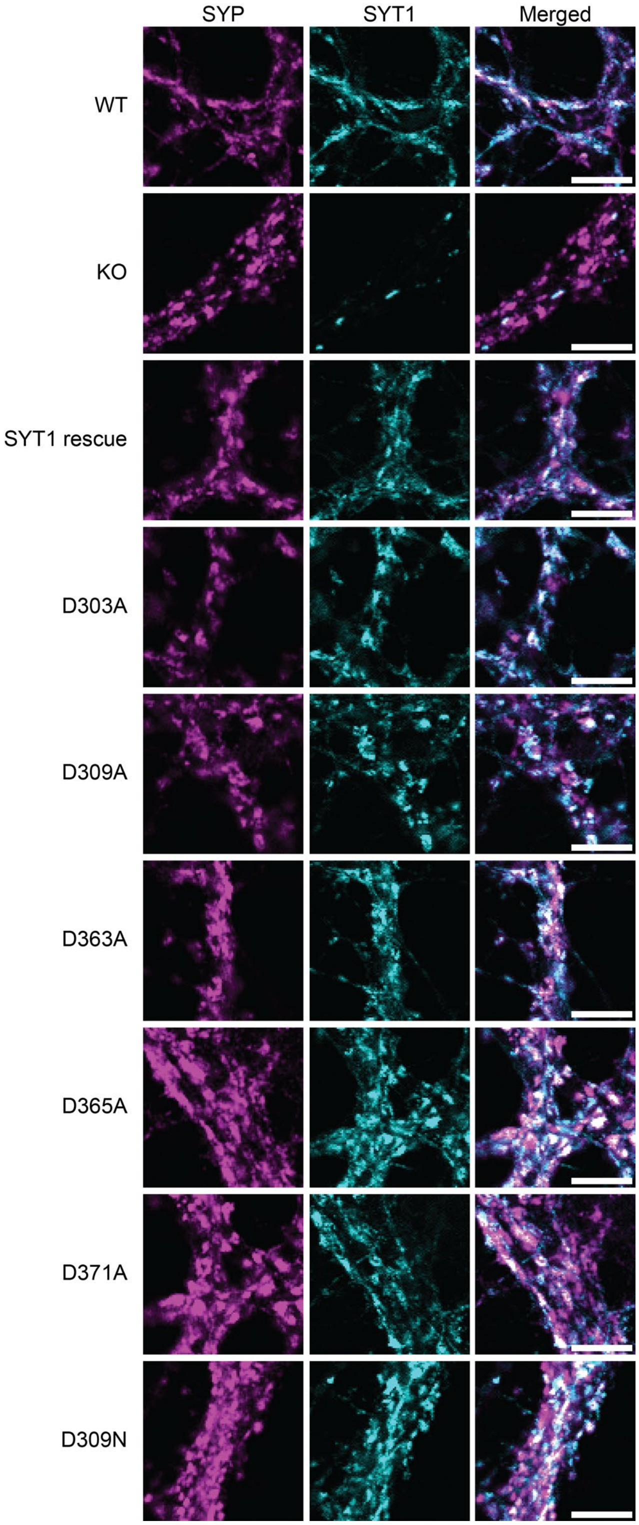


**Fig. S6: C2B $Ca^{2+}$ ligand mutants are localized to SVs.** Representative images showing co-localization of C2B CLMs with the SV marker synaptophysin (SYP). Scale bar, 5 μm.

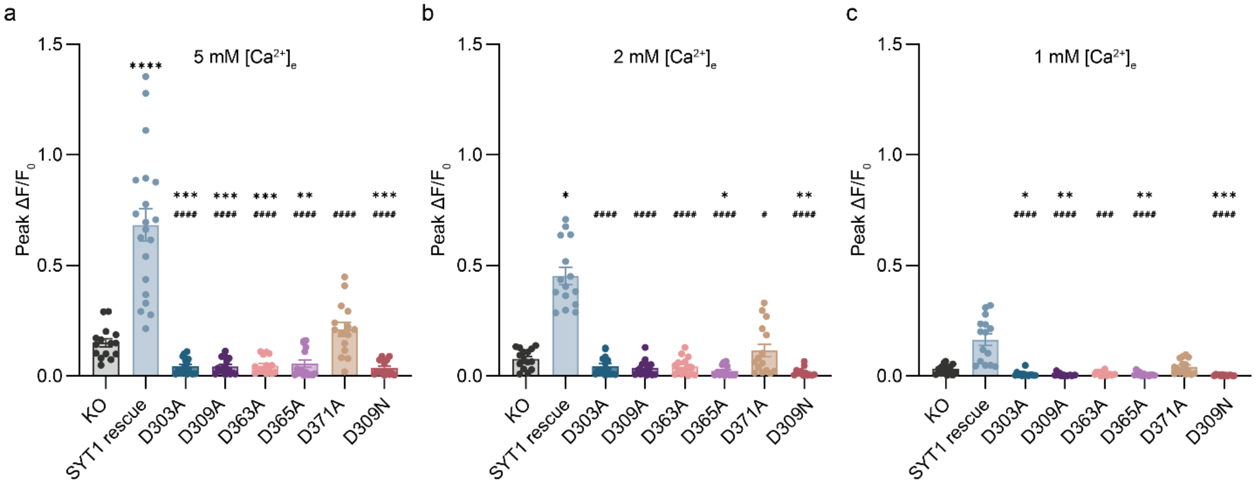


**Fig. S7: C2B CLMs disrupt glutamate release at all $[Ca^{2+}]_e$:** Peak iGluSnFR $\Delta F/F_0$ of C2B CLMs at 5, 2, 1 μM $[Ca^{2+}]_e$. Each data point represents a single FOV, n = at least 14 FOVs, from at least 3 independent trials; error bars indicate SEM. Statistical significance was determined using the Brown-Forsythe ANOVA test, with Dunnett's T3 multiple comparisons test. At lower $[Ca^{2+}]_e$, D371A fails to yield release, and other mutants perform worse than *syt1* KOs. For all panels, ****, #### $p < 0.0001$; ***, ### $p < 0.001$; ** $p < 0.01$; *,# $p < 0.05$.

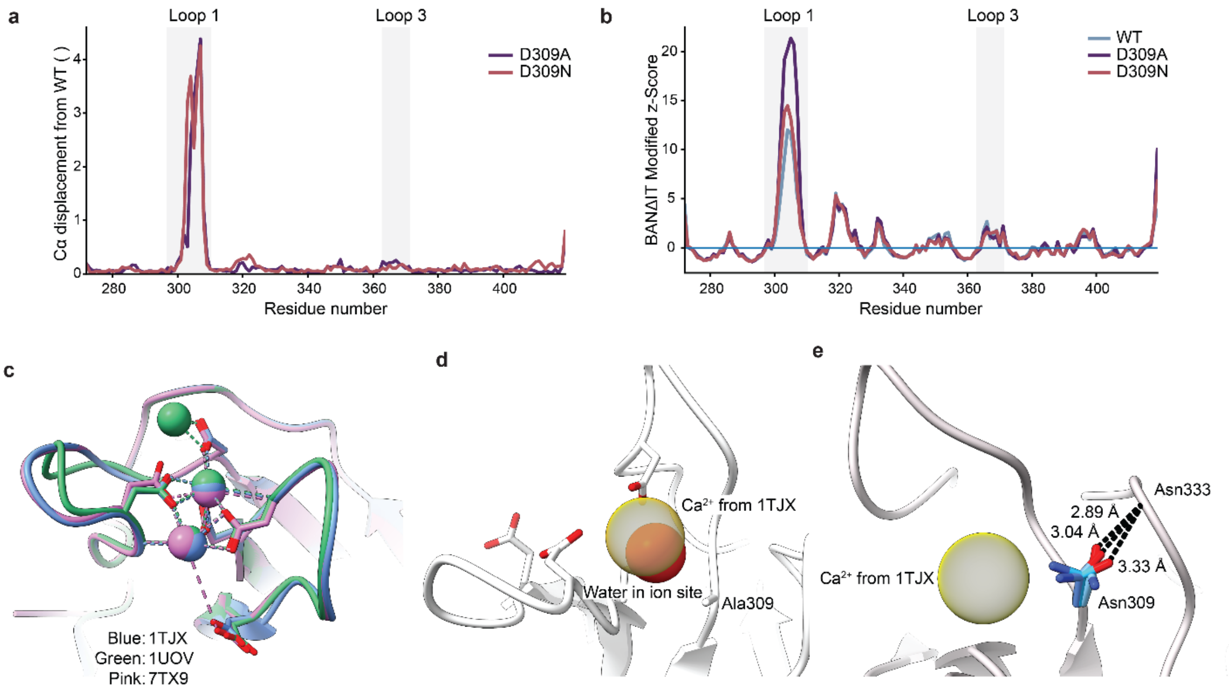


**Fig. S8: Loop 1 displacement and normalized mobility in WT and pathogenic C2B mutants. a,** Per-residue Cα displacement of D309A and D309N relative to WT SYT1 C2B following alignment of the conserved β-sandwich core. The largest modeled differences are localized to loop 1 (K297–P310), whereas loop 3 (D363–D371) remains largely unchanged. **b,** BANΔIT Modified z-Score profiles calculated from Cα B-factors for WT, D309A, and D309N. Shaded regions indicate loop 1 and loop 3. Loop 1 shows elevated normalized B-factors in all three structures, with the largest values in D309A. **c,** Overlay of the $Ca^{2+}$-bound SYT1 C2B structures 1TJX, 1UOV, and 7TX9 following alignment of the conserved β-sandwich core. The five acidic $Ca^{2+}$-binding residues and all deposited $Ca^{2+}$ ions are shown; D309 directly coordinates $Ca^{2+}$ across the three reference structures. Exact D309–$Ca^{2+}$ distances are listed in Table S3. **d,** D309A structure with the principal D309-coordinated $Ca^{2+}$ position from 1TJX was transferred into the mutant coordinate frame. A modeled water occupies the same solvent-filled region, indicating that removal of the D309 carboxylate does not occlude the binding site. **e,** D309N showing the three modeled Asn309 conformations and their interactions with the N333 backbone. The principal D309-coordinated $Ca^{2+}$ position from 1TJX is shown as a transparent reference sphere. The observed Asn309 carbonyl positions do not reproduce the principal WT $Ca^{2+}$-coordination geometry. Normalized Cα B-factors were computed using BANΔIT[4].

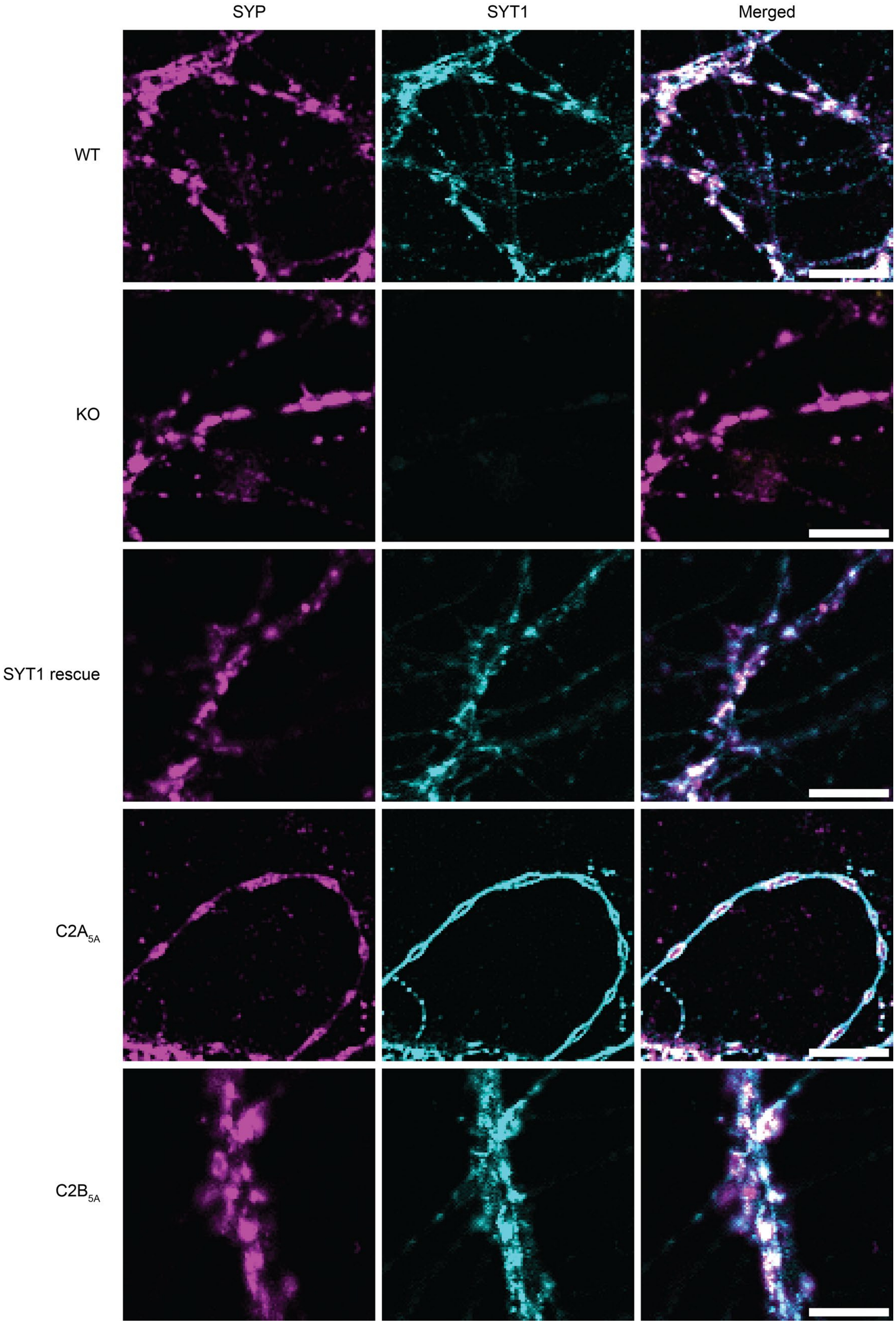
SYP
SYT1
Merged
WT
KO
SYT1 rescue
C2A5A
C2B5A

**Fig. S9: C2A$_{5A}$ is localized to the axonal plasma membrane, whereas C2B$_{5A}$ is localized on SVs.** Representative images showing co-localization of C2B CLMs with the SV synaptophysin (SYP). Scale bar, 5 µm.

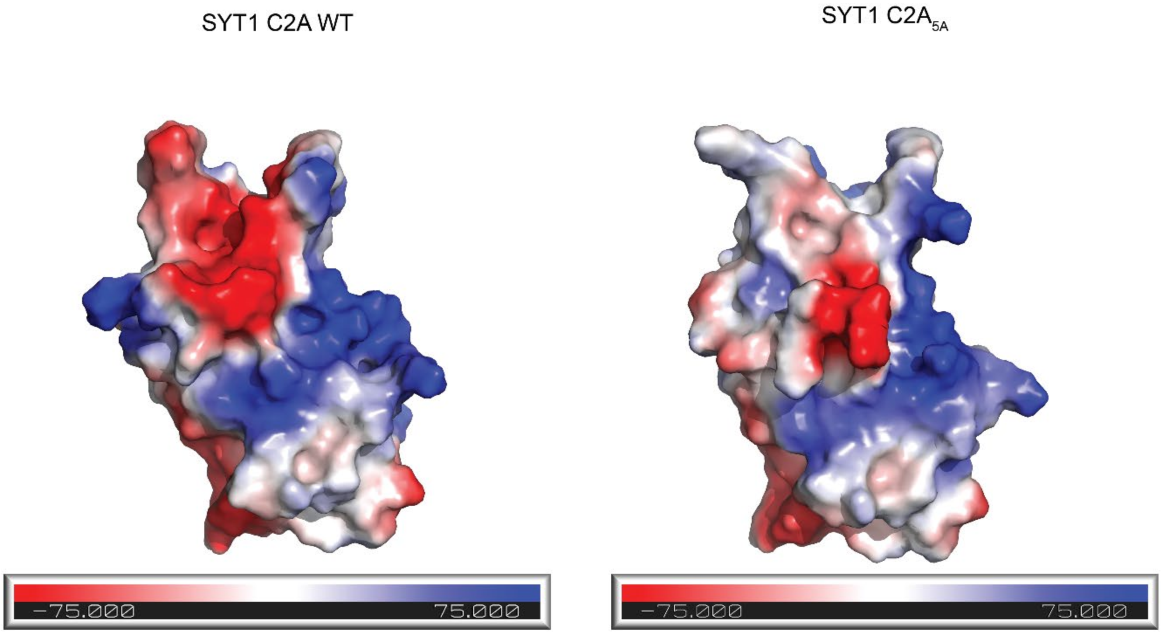


**Fig. S10: Ca$^{2+}$ sensing loops in C2A$_{5A}$ are neutralized.** Electrostatic surface potential of WT C2A and C2A$_{5A}$ showing neutralization of all negative charge in the Ca$^{2+}$-sensing loops. Red, blue, and white in the color ramp depict negative, positive, and neutral charges, respectively. C2A$_{5A}$ model was generated using AlphaFold[5]. The electrostatic surface was rendered using PyMOL[3].
